# Colorectal cancers with distinct metastatic potential trigger divergent early T cell responses

**DOI:** 10.64898/2026.06.30.735606

**Authors:** Marwa Saad, Alexandra Thoms, Yue S. Yin, Izabela Mamede, Afsana Rahman, Ruonan Chen, Sebastián E. Carrasco, Patrick W. Darcy, Norihiro Goto, Sohail F. Tavazoie, Angelina M. Bilate, Daniel Mucida

## Abstract

Colorectal cancer (CRC) remains a leading cause of cancer mortality, with most cases refractory to immunotherapy. Distinguishing tumor-induced from steady-state mucosal T cell responses has been a critical barrier to understanding antitumor immunity in CRC. Using orthotopic transplantation of CRC organoids with and without metastatic potential, combined with temporal T cell fate-mapping, we show that non-metastatic tumors elicit early recruitment of CD8αβ⁺ and CD4⁺ T cells that acquired cytotoxic and Th1-like programs, whereas pro-metastatic tumors induce a naïve-like, hypoactivated state. Tumor-infiltrating CD4^+^ T cells underwent clonal expansion, including clones recognizing microbial and dietary antigens. T cells in physical contact with tumor cells, identified by uLIPSTIC, were enriched for expanded and cytotoxic clones. Fate-mapped T cells from non-metastatic tumors suppressed tumor growth in an IFN-γ-dependent manner, whereas pro-metastatic tumor-derived T cells failed to do so. Mechanistically, pro-metastatic tumors downregulated MHCII, and *Ciita* targeting in non-metastatic organoids reduced CD4⁺ clonal expansion and led to tumor progression. Together, these findings define divergent early T cell trajectories associated with CRC metastatic potential, indicating that ineffective local immune engagement precedes metastatic dissemination.

## Introduction

The intestinal mucosa harbors the largest immune compartment in the mammalian body, where resident and recruited immune cells must maintain tolerance to commensal and dietary antigens while preserving their ability to mount effective responses against pathogens and malignant transformation^1,2^. Uncontrolled epithelial proliferation leads to colorectal cancer (CRC), the second leading cause of cancer-related deaths worldwide^3–5^. More than 85% of human CRC cases arise from accumulating sporadic or hereditary mutations in the tumor suppressor genes *Apc, Tp53, Smad4*, and the oncogene *Kras*^3,6,7^. Although immune checkpoint blockade (ICB) has transformed the treatment of certain solid tumors, most CRCs, particularly those with a proficient mismatch repair system (pMMR), remain refractory to immunotherapy and are prone to metastasis^8–10^. In contrast, mismatch repair-deficient (dMMR) tumors harbor a high mutational burden that promotes immunogenicity and responsiveness to T cell-mediated therapies^9,10^. This dichotomy underscores the pivotal yet incompletely understood role of local T cell responses in shaping CRC progression and its metastatic propensity^11^.

Although adaptive immunity can eliminate nascent tumor cells, it can also become functionally silenced or co-opted in an evolving tumor microenvironment (TME)^12–14^. In the gut tissue, this challenge is compounded by the constant need to restrain damaging immune activation toward abundant luminal antigens, creating an environment in which tumor-specific responses must arise amid pre-existing tolerance mechanisms^15–18^. However, how T cells integrate these conflicting cues, especially during the early stages of tumor growth and metastatic dissemination, remains unclear^11^. Previous studies have largely focused on either highly immunogenic dMMR CRC models or non-physiological subcutaneous transplants, providing limited insight into gut-resident or metastasis-associated T cell responses^19,20^. Additionally, polyclonal T cell responses in non-metastatic versus pro-metastatic CRC, distinct from “background” gut immune responses, remain unexplored.

To address these gaps, we combined orthotopic CRC models with distinct metastatic potentials and a temporal T cell fate-mapping strategy that enabled discrimination between pre-existing intestinal T cells and newly recruited potential antitumor clones. This approach allowed us to compare the recruitment, activation, and clonality of T cells in non-metastatic and pro-metastatic tumors within their native microenvironments. We found that these tumor types elicit fundamentally distinct early T cell activation responses, ranging from robust effector differentiation and clonal expansion to naïve-like or hypoactivated states, highlighting a previously unrecognized divergence in local immune engagement that precedes metastatic progression.

## Results

### Immune profiling of orthotopic CRC models with distinct metastatic potential

To investigate T lymphocyte infiltration in colorectal tumors and adjacent healthy colon epithelium, we implemented colonoscope-guided orthotopic models of CRC^21,22^. We injected the following mouse tumor organoid lines in the colonic walls of immunocompetent B6 mice: 1) AKP, harboring *Apc^−/–^*, *Kras^G12D^*, and *Tp53^−/–^ mutations*^21,23,24^, and 2) pro-Metastatic (pro-Met) line which was generated by multiple *in vivo* passaging of the AKP line (**Figures 1A, S1A and B**). For each of these transplanted tumor organoid lines, we assessed colon tumor growth, liver metastasis, and T cell phenotype at an early stage (two weeks post-tumor injection) and a late stage (6-7 weeks after tumor injection) of CRC development. This temporal analysis enabled us to determine when differences in tumor aggressiveness and immune infiltration first emerge and how they evolve over time. While the average colon tumor diameter was comparable between both tumor organoid models at the early CRC stage, pro-Met tumors were larger at the late CRC stage (**Figure 1B**). Liver macrometastases were observed only in the pro-Met model at the late CRC stage (**Figures 1C and S1C**). These observations establish AKP and pro-Met as matched primary CRC models that diverge sharply in metastatic behavior, providing a framework for further investigation of their primary TME.

**Figure 1:**
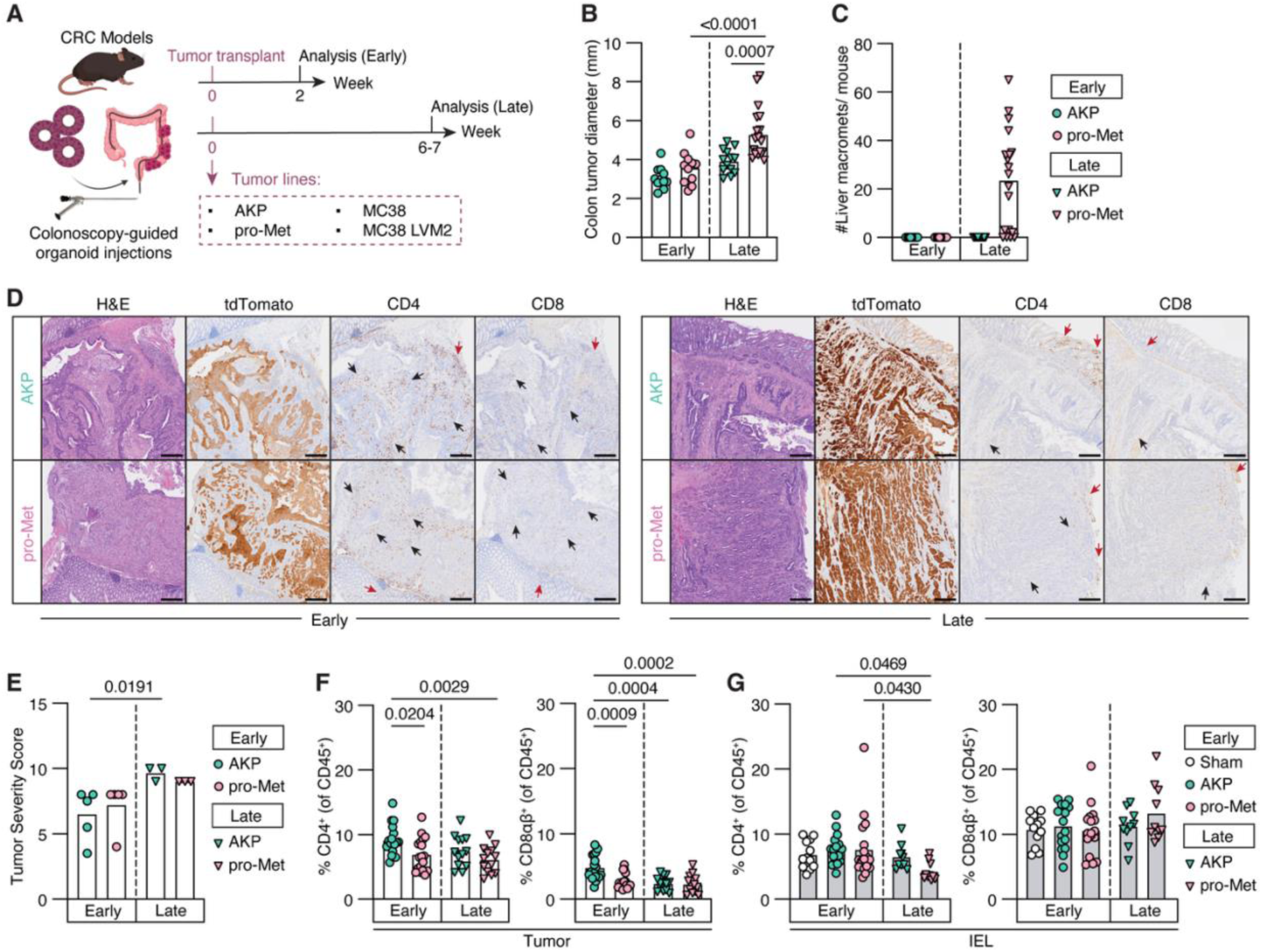
Characterization of tumor and T cell phenotypes in orthotopic CRC models with distinct metastatic potential. (A) Experimental design. Colonoscopy-guided injection of various tumor lines in the mouse colon wall. Mice were harvested for analysis at 2 weeks (early) or 6-7 weeks (late) post-tumor injection. (B) Colon tumor diameter and (C) number of liver macrometastases per mouse in the early and late stages of the indicated tumor organoid lines. Each symbol represents a mouse. (D) Hematoxylin and Eosin (H&E) staining and immunohistochemistry for tdTomato (demarcating tumor cells), CD4, and CD8α in AKP and pro-Met tumor sections at the early (left) and late (right) CRC stages. Black arrows point to T cells inside tumor bed, and red arrows point to T cells at the tumor periphery. Scale bar = 250 μm. (E) Tumor severity score (based on necrosis, fibroplasia, and epithelial dysplasia grades) of colonic AKP and pro-Met tumors assessed at the early and late CRC stages. Each symbol represents a mouse. (F) Frequencies of CD4^+^ and CD8αβ^+^ T cells among CD45^+^ T cells in colon tumors of respective tumor lines at the early and late CRC stages. Each symbol represents a mouse. (G) Frequencies of CD4^+^ and CD8αβ^+^ T cells among CD45^+^ IEL isolated from colon epithelium of sham-or tumor-injected mice at the early and late CRC stages. Each symbol represents a mouse. Ordinary one-way analysis of variance (ANOVA) with Tukey’s multiple comparison test was performed for (B), (C), (F), and (G); and Kruskal-Wallis test followed by Dunn’s multiple comparison test in (E).

To determine how tumor morphology and immune infiltration differ across models and time points, we performed histopathological analysis of colons of mice bearing AKP and pro-Met tumors at both early and late stages. We performed hematoxylin and eosin (H&E) staining combined with immunohistochemistry (IHC) for tdTomato (to demarcate tumor cells) and CD4 and CD8 to assess T cell infiltration in colon lesions (**Figure 1D**). Both tumor types were consistent with adenocarcinoma in most of the analyzed tumors, with invasion extending into the muscularis at an early stage (**Figure 1D**). In the late stage, both models were uniformly classified as adenocarcinomas and exhibited transmural extension from the mucosa to the serosa (**Figures 1D and S1D**). To quantify tumor progression, we used a tumor severity score by integrating necrosis, fibroplasia, and epithelial dysplasia grades. Using this metric, AKP tumors exhibited an increase in severity as they progressed from early to late stages, reflecting progressive histopathological worsening over time. In contrast, pro-Met tumors did not exhibit a significant stage-dependent change, consistent with the presence of a more advanced histopathological state early on. Early pro-Met tumors scored intermediate, which was not significantly different from either early AKP tumors or late pro-Met tumors, whereas late AKP and late pro-Met tumors were indistinguishable (**Figure 1E**). IHC assessment indicated that both tumors were highly infiltrated with CD4^+^ T cells and, to a lesser extent, with CD8^+^ T cells in the early stage. However, in the late stage, CD4^+^ and CD8^+^ T cells were mostly detected at the tumor periphery, with less infiltration inside the tumor bed (**Figure 1D**). To quantitatively assess these immune differences, we performed flow cytometry on colon tumor tissues to measure T cell accumulation across models and time points. We observed higher CD4^+^ and CD8αβ^+^ T cell accumulation in early stage AKP tumors than in pro-Met tumors at both early and late stages (**Figure 1F**). In addition to colon tumors, we examined adjacent non-tumor epithelium (for intraepithelial lymphocytes, referred to as IEL) to serve as a reference baseline and to assess whether tumors alter adjacent non-tumor regions. Colonic IEL from sham-injected mice were used as controls to account for the potential damage caused by colonoscopic injection. The non-tumor epithelium of AKP-and pro-Met-injected mice showed CD4^+^ and CD8αβ^+^ IEL accumulation comparable to that of sham-injected controls (**Figure 1G**). These data suggest that the AKP model establishes a more inflamed tumor environment than the pro-Met model in the early stages of CRC development.

To complement the organoid models and include a widely used dMMR CRC model, we orthotopically injected the murine colon adenocarcinoma cell line MC38 and its derivative (MC38 LVM2), which was passaged twice in the mouse liver (**Figure 1A**). In the early stage, mice transplanted with MC38 or MC38 LVM2 cells developed colon tumors comparable in size to late-stage pro-Met tumors and significantly larger than early stage AKP tumors (p = 0.0352 for MC38 vs. AKP; p = 0.0052 for MC38 LVM2 vs. AKP) (**Figure S1E**). Despite this, no liver metastases were detected in the MC38 cell lines. Because mice bearing MC38 tumors exhibited morbidity beyond the early time point, all subsequent analyses of this model were conducted exclusively 2 weeks post-injection. Flow cytometry analysis revealed that MC38 and MC38 LVM2 tumors harbored comparable frequencies of CD4⁺ and CD8αβ⁺ T cells. Although MC38 represents a dMMR CRC model, CD4⁺ T cell accumulation in MC38 tumors was lower than that observed in early stage AKP tumors (p = 0.0042 for MC38; p = 0.0057 for MC38 LVM2) (**Figure S1F**). In the non-tumor epithelium, CD4⁺ and CD8αβ⁺ IEL frequencies in MC38-injected mice were similar to those in the MC38 LVM2- and sham-injected controls (**Figure S1G**). These results suggest that although MC38 and MC38 LVM2 tumors grow more aggressively than the organoid models tested, they provide a dMMR orthotopic CRC system suitable for analyzing T cell responses at the tumor site. Collectively, these data indicate distinct T cell responses in orthotopic CRC models with divergent growth patterns, microsatellite stability, and metastatic potential.

### Non-metastatic AKP tumors induce potent early T cell responses

The early-stage divergence in CD4⁺ and CD8αβ⁺ T cell accumulation between AKP and pro-Met tumors suggests that these models establish distinct immune microenvironments at the tumor onset. This led us to pursue two complementary approaches to further investigate these differences. First, we evaluated the potential of T cells to directly interact with AKP and pro-Met tumor cells at the early CRC stage, given that IHC confirmed their close proximity to both tumor types at this time point (**Figure 1D**). To achieve this, we first adapted the universal LIPSTIC (uLIPSTIC) system to our CRC models^25,26^. Sortase A (SrtA)-expressing AKP and pro-Met organoid lines were orthotopically injected into the colon walls of G5-expressing *Pou2f3*^CreERT2^ uLIPSTIC mice (**Figures 2A and S1H**). Treatment of these mice with tamoxifen induced SrtA expression in tuft cells and thymic epithelial cells, thus preventing rejection of SrtA-expressing tumors (**Figure 2A**)^27^. Two weeks after tumor injection, the mice were injected with biotin-LPETG substrate, and the tumors were harvested for flow cytometry analysis. AKP and pro-Met tumors expressed SrtA, FLAG tag, and biotin on their surface, confirming their ability to transfer biotin-LPETG substrate to G5-expressing T cells upon intercellular interaction (**Figure 2B**). While pro-Met tumors showed minimal interactions with CD8αβ^+^ and no detectable interaction with CD4^+^ T cells, a sizable fraction of CD4^+^ and CD8αβ^+^ T cells interacted with AKP tumors (**Figure 2C**). Compared to non-interacting (biotin^−^) T cells, a higher proportion of biotin^+^ T cells expressed the activation markers PD-1, Tbet, and CD44, suggesting that the interactions between T cells and AKP tumors are biologically functional (**Figure 2D**). Colonic tuft cells did not label any immune cells within the colonic IEL, lamina propria (LP), or mesenteric lymph nodes (mLN), further supporting that any observed labeling in the tumor originates from interactions with tumor cells expressing SrtA (**Figures S1I and J**). To determine if αβ T cells are the primary immune interactors in this model, we profiled all tumor-infiltrating CD45^+^ populations, including B cells and various myeloid lineages. We found that αβ T cells constitute the vast majority (∼70%) of the biotin^+^ interacting fraction, while innate populations such as eosinophils, neutrophils, and macrophages showed minimal biotin labeling (**Figure S1K**). This reveals that despite T cells representing a minority of the total CD45^+^ infiltrate, they are the most interactive immune population with the tumor cells. These findings reveal that early AKP tumors support productive T cell engagement, whereas pro-Met tumors largely evade such interactions, establishing divergent immune trajectories between the two tumor types.

**Figure 2:**
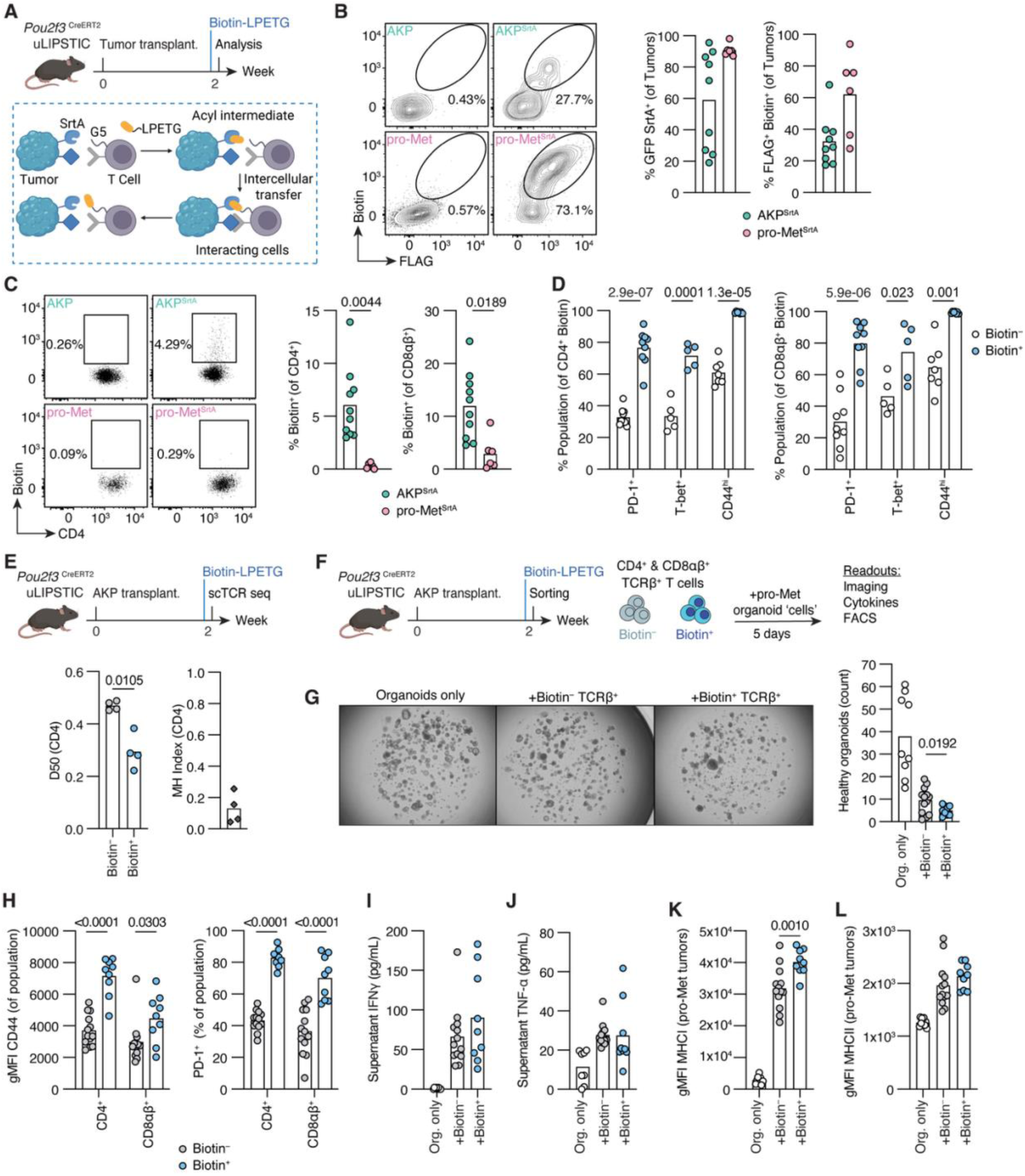
Non-metastatic AKP tumors but not pro-Met functionally interact with T cells. (A) Experimental design. *Pou2f3*^CreERT2^ uLIPSTIC mice were orthotopically injected with AKP^SrtA^ or pro-Met^SrtA^ organoids (or respective control organoids lacking SrtA expression) using a colonoscope. Two weeks later, six doses of biotin-LPETG substrate were injected intraperitoneally over a course of 2 h (8 µmol/1^st^ dose, followed by 2 µmol for each subsequent dose). Tumors and tumor-infiltrating T cells were harvested for flow cytometry analysis 1 h after the last substrate injection. (B) Dot plots of harvested tumor cells showing the surface expression of FLAG and Biotin. Frequencies of GFP SrtA^+^ (middle) and FLAG^+^ Biotin^+^ (right) among all tumor cells. (C) Dot plots showing biotin expression in CD4^+^ T cells harvested from the indicated tumors, and frequencies of biotin^+^ among CD4^+^ (middle) and CD8αβ^+^ (right) T cells harvested from the indicated tumors. (D) Frequencies of indicated populations (PD-1^+^, T-bet^+^, CD44^hi^) among biotin^−^ or biotin^+^ CD4^+^ (left) or CD8αβ^+^ (right) T cells. (E) scTCR seq of biotin^−^ and biotin^+^ CD4^+^ T cells isolated from mice bearing AKP^SrtA^ tumors at the early stage following biotin-LPETG administration as described in A. Diversity index 50 (D50) of CD4^+^ T cell clones (left), and Morisita-Horn (MH) index showing overlap between biotin^−^ and biotin^+^ CD4^+^ T cell clones. (F) Experimental design. *Pou2f3*^CreERT2^ uLIPSTIC mice were orthotopically injected with AKP^SrtA^ organoids and injected with biotin-LPETG substrate as indicated. Biotin^−^ and biotin^+^ CD4^+^ and CD8αβ^+^ T cells were bulk sorted and pooled from n=3-4 mice per experiment. Sorted T cells were then co-cultured *in vitro* with pre-dissociated pro-Met tumor organoids for five days (4000 T cells and 1000 pro-Met organoid cells per well). Data is pooled from 3 independent experiments. (G) Brightfield (BF) imaging (2X, 1.5 Zoom) of control (organoids only), in addition to organoids co-cultured with biotin^−^ and biotin^+^ T cells. The number of healthy organoids was quantified (right). Healthy organoids were defined as those with well-defined edges with limited to no debris inside the organoid. (H) Geometric mean fluorescence intensity (gMFI) of CD44 (left), and frequency of PD-1^+^ CD4^+^ and CD8αβ^+^ T cells among biotin^−^ and biotin^+^ T cells isolated 5 days after co-culture. (I–J) Cytometric bead array (CBA) analysis of co-culture supernatant collected on day 5. Graphs show levels of IFN-γ (I) and TNF-α (J) per condition. (K–L) gMFI of MHCI (K) and MHCII (L) expression assessed by flow cytometry on pro-Met tumor organoids harvested at day 5. Unpaired Student’s t-tests were performed and p-values are indicated. (B-E), each symbol represents a mouse. (G-L), each symbol represents a co-culture well.

To determine whether the physical interactions captured by uLIPSTIC identify a functionally relevant population, we first sorted tumor-infiltrating biotin^+^ and biotin^−^ CD4^+^ T cells from AKP-bearing mice and performed single-cell TCR sequencing (scTCR-seq) (**Figure 2E**). We estimated clonal diversity with the diversity 50 index (D50) of TCRβ, with values ranging from 0 (least diverse) to 0.5 (most diverse)^28^. We also used the Morisita-Horn Index (MHI) to determine clonal overlap between the biotin^+^ and biotin^−^compartments, with 1 indicating 100% clonal overlap. The biotin^+^ CD4^+^ T cells exhibited a significantly lower D50 compared to the biotin^−^ fraction, demonstrating that physical engagement with the tumor is associated with clonal expansion ( **and S1L**). Additionally, minimal clonal overlap was observed between the biotin^+^ and biotin^−^ compartments, as indicated by the low MHI (**Figure 2E**). Secondly, to assess the functional potency of the interacting repertoire, we co-cultured biotin^+^ and biotin^−^ αβ T cells with pro-Met organoids (**Figure 2F**). Both biotin^+^ and biotin^−^ αβ T cells exhibited killing capacity, with the biotin^+^ fraction significantly reducing organoid survival compared to the biotin^−^ fraction (**Figure 2G and S1M**). This cytotoxic activity was accompanied by an activated CD44^hi^ PD-1^+^ phenotype (**Figure 2H**). Both biotin^+^ and biotin^−^ T cell fractions induced robust cytokine production relative to organoids cultured alone, as shown by increased IFN-γ and TNF-α levels in the co-culture supernatants. Although cytokine secretion was not significantly different between the biotin^+^ and biotin^−^ conditions (**Figures 2I and J**), biotin^+^ T cells induced a greater upregulation of MHC class I (MHCI) expression on the pro-Met tumor organoids (**Figure 2K**). MHC class II (MHCII) expression was similarly elevated in both T cell-containing conditions compared with organoids alone, with no significant difference observed between the biotin^+^ and biotin^−^fractions (**Figure 2L**). Given that MHCI expression is often more sensitive to low-level cytokine exposure and contact-dependent signals than MHCII^29^, the greater MHCI upregulation by the biotin^+^ fraction may indicate a stronger effector interaction with tumor cells. These findings indicate that T cells physically interacting with AKP tumors *in vivo* constitute a clonally expanded, activated, and functionally cytotoxic population with enhanced tumor-modulating capacity.

### A fate-mapping system to identify tumor-induced T cells in orthotopic CRC

The orthotopic CRC-uLIPSTIC system is a powerful tool for capturing direct cellular interactions during the short window of substrate administration. However, a significant roadblock in characterizing adaptive immune responses to tumors in polyclonal systems is the inability to distinguish recently recruited T cells responding to tumors (or tumor-related perturbations) from tissue-resident T cells that occupy the transplantation site before tumor introduction. This distinction is particularly challenging in the intestinal environment, given the vast number of resident lymphocytes that participate in ongoing immune responses against luminal antigens^30,31^. To address this challenge and complement the uLIPSTIC system, we combined orthotopic CRC models with a fate-mapping approach to track peripheral T cell recruitment and activation in the colon^21,22,32,33^. Tamoxifen treatment of *Sell*^CreERT2^ X *Rosa26*^CAG-LSL-tdTomato^ (i*Sell*^Tomato^) mice permanently labels naïve T cells expressing L-selectin (*Sell*, CD62L^+^) as Tomato^+^. This strategy allowed us to enrich for “ex-naïve” (CD62L^−^ Tomato^+^) T cells and follow the fate of tumor-induced T cells^32,33^.

To validate the T cell labeling efficiency in i*Sell*^Tomato^ mice under homeostatic conditions, we harvested the mLN, colon epithelium, and LP shortly after tamoxifen administration for flow cytometry analysis (**Figures S2A and B**). Across all compartments, more than half of the CD62L^+^ cells were Tomato^+^. As expected, the majority of CD4^+^ and CD8αβ^+^ T cells in the mLN expressed CD62L. In contrast, approximately 20–40% of CD4^+^ and CD8αβ^+^ T cells in the colon epithelium and LP were CD62L^+^, mainly due to the labeling of naïve T cells naturally present in isolated lymphoid follicles and colonic patches^34^. Immunofluorescence (IF) staining and confocal imaging confirmed the presence of Tomato^+^ cells co-localizing with lymphoid aggregate structures in the colon (**Figure S2B**). To further validate that i*Sell*^Tomato^system could also be used to label and track T cells during established tumor growth, mice were treated with tamoxifen four weeks after AKP or pro-Met tumor transplantation and analyzed one day after (**Figure S2C**). IF staining confirmed the presence of CD3^+^ Tomato^+^ T cells within tumor-bearing colons, with these cells predominantly localized to lymphoid aggregates (**Figure S2D**). These findings demonstrate that i*Sell*^Tomato^ labeling remains effective at later stages of tumor progression and can be used to track newly recruited T cells in the tumor-bearing colon. Throughout this study, the term “fate-mapped” refers specifically to these CD62L^−^ Tomato^+^ T cells. Because CD62L downregulation can occur following T cell activation in multiple anatomical locations, this population likely includes cells primed in tumor-draining lymph nodes, colonic lymphoid structures, or within the TME itself. Thus, the i*Sell*^Tomato^ system is not intended to identify exclusively tumor-specific T cells, but rather to enrich for the wave of polyclonal T cells that are recruited and activated during the defined period of tumor establishment, enabling their distinction from pre-existing tissue-resident lymphocytes.

To investigate T cell recruitment and activation dynamics in orthotopic CRC models, we injected the previously described tumor organoids and cell lines into the colonic walls of i*Sell*^Tomato^mice and administered tamoxifen either before or four weeks after tumor injection. Two weeks after tamoxifen administration, we collected colon tumors and epithelium from tumor- and sham-injected mice for flow cytometry analysis (**Figure 3A**). In the early stage, we observed enhanced accumulation of newly recruited, fate-mapped CD62L^−^ Tomato^+^ CD4^+^ and CD8αβ^+^ T cells in non-metastatic AKP compared to pro-Met colon tumors (**Figures 3B and C**). In the late stage, CD4^+^ and CD8αβ^+^ T cell recruitment to non-metastatic AKP tumors decreased to levels comparable to those observed in both stages of the pro-Met model (**Figures 3B and C**). Both MC38 and MC38 LVM2 tumors displayed T cell recruitment rates comparable to those of early stage pMMR AKP tumors (**Figure 3D**). The early stage AKP model resulted in increased accumulation of newly recruited CD4^+^ T cells in the adjacent colon epithelium, whereas other models displayed recruitment patterns comparable to the sham control (**Figure 3E**). In contrast, neither the organoid nor cell line models affected the accumulation of newly recruited CD8 αβ^+^ T cells in the adjacent colon epithelium (**Figures 3F and G**). These data further indicate that non-metastatic AKP tumors trigger new T cell accumulation and foster a more inflamed microenvironment, in contrast to pro-Met tumors, which appear to evade such new responses from the early stage, eliciting a T cell response similar to that in late-stage CRC.

**Figure 3:**
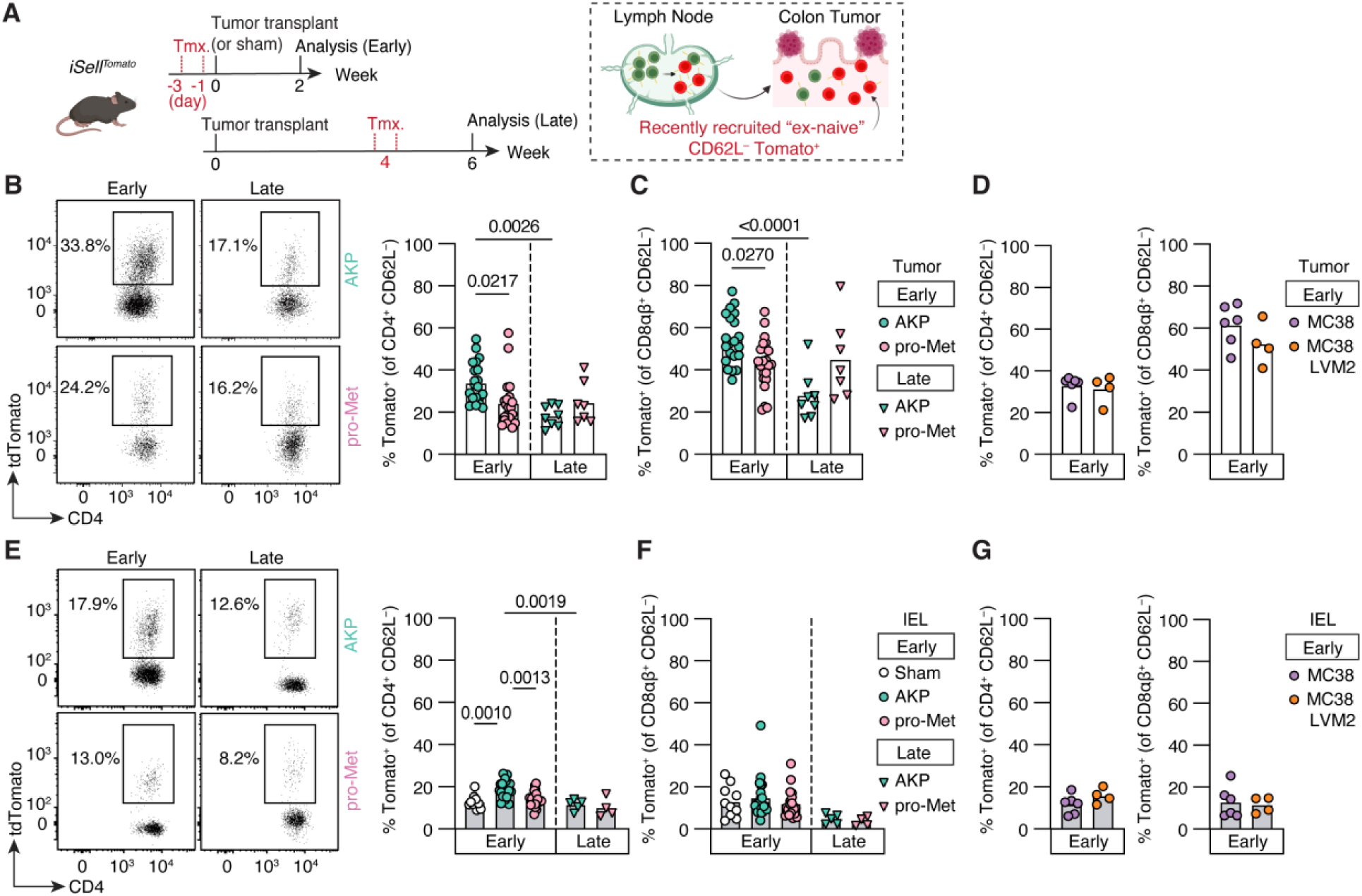
Non-metastatic AKP tumors induce potent antitumor early T cell responses. (A) Experimental design. i*Sell*^Tomato^ mice were treated with two tamoxifen doses either before or four weeks after orthotopic tumor injection. Sham injection was performed as control. Flow cytometry analyses of T cells were performed two weeks after tamoxifen administration. (B-C) Dot plots and frequencies of recently recruited Tomato^+^ T cells among CD4^+^ CD62L^−^ (B) and CD8αβ^+^ CD62L^−^ (C) T cells in AKP and pro-Met colon tumors at the early and late CRC stages. (D) Frequencies of recently recruited Tomato^+^ T cells among CD4^+^ CD62L^−^ (left) and CD8αβ^+^ CD62L^−^(right) T cells in MC38 and MC38 LVM2 colon tumors at the early CRC stage. (E-F) Dot plots and frequencies of recently recruited Tomato^+^ T cells among CD4^+^ CD62L^−^ (E) and CD8αβ^+^ CD62L^−^ (F) IEL isolated from colon epithelium of sham- or tumor-injected mice at the early and late CRC stages. (G) Frequencies of recently recruited Tomato^+^ T cells among CD4^+^ CD62L^−^ (left) and CD8αβ^+^ CD62L^−^(right) IEL isolated from colon epithelium of tumor-injected mice at the early CRC stage. Unpaired Student’s t-tests were performed for (D) and (G). Ordinary one-way ANOVA with Tukey’s multiple comparison tests were performed in (B), (C), (E), and (F). (B-G), each symbol represents a mouse.

### Impaired early T cell response is not sufficient to drive metastasis of AKP tumors

The early T cell phenotypic differences between the AKP and pro-Met models prompted us to investigate whether the more inflamed T cell environment in AKP tumors could impact their metastatic progression. First, we determined whether the absence of metastasis after AKP tumor transplantation reflects an intrinsic inability of AKP cells to colonize the liver. Upon intrasplenic injection^35,36^, both AKP and pro-Met tumor lines were equally capable of forming liver metastases, indicating that AKP tumors are competent to grow in the liver when given direct access, and that their non-metastatic outcome in the orthotopic model is likely due to impaired dissemination rather than inability to colonize the liver (**Figure S2E**).

We then directly investigated the role of T cells in shaping the AKP primary tumor and metastatic outcomes. We orthotopically injected AKP and pro-Met tumors into immunodeficient *Rag1*^−/–^ (RKO) mice, which lack functional T and B cells. By six weeks post-injection, AKP and pro-Met colon tumors in RKO mice reached sizes comparable to those observed in immunocompetent hosts. Consistent with their phenotypes in immunocompetent mice, pro-Met tumors formed liver metastases, whereas AKP tumors remained nonmetastatic (**Figure S2F**). To specifically test the impact of T cells on the observed AKP phenotype, we performed antibody depletion of CD4^+^ or CD8^+^ T cells in C57BL/6 mice transplanted with AKP tumor organoids (**Figure S2G**). Efficient depletion of CD4^+^ or CD8^+^ T cells was confirmed by flow cytometry (**Figure S2H**). At the early time point, AKP tumor sizes were comparable across the groups. However, at the late stage, CD8-depleted mice developed significantly larger tumors, whereas CD4^+^ T cell-depleted mice showed intermediate tumor sizes between the isotype control and CD8^+^ T cell-depleted conditions (**Figure S2I**). As expected, CD4^+^ T cell depletion impaired CD8^+^ T cell activation, reducing their IFN-γ production and PD-1 expression, whereas CD8 depletion had minimal effects on CD4^+^ T cell activation (**Figures S2J and K**). Nonetheless, liver metastases were not detected in any of the experimental groups, suggesting that the intrinsic transcriptional programs in AKP and pro-Met tumors likely play a dominant role in determining their metastatic fate. Although neither CD4⁺ nor CD8⁺ T cell targeting was sufficient to drive metastatic progression in AKP tumors, our data reinforce the role of T cells in shaping the primary TME. This prompted us to investigate how CD4⁺ and CD8αβ⁺ T cells functionally adapt to distinct AKP versus pro-Met TMEs, independent of their capacity to determine metastatic outcomes.

### Transcriptional profiling of recently recruited T cells at early stage non-metastatic and pro-metastatic CRC

To resolve how early T cell adaptation programs diverge in AKP versus pro-Met CRC models, we focused on the fate-mapping i*Sell*^Tomato^ approach, which enables tracking of polyclonal “ex-naïve” T cells labeled at the time of tumor transplantation and during the earliest stages of tumor development. This approach allowed us to assess how newly recruited T cells adapt to the TME. i*Sell*^Tomato^ mice were transplanted with AKP or pro-Met organoids, or subjected to a sham procedure (*see* **Figure 3A**). Two weeks later, we sorted CD62L^−^ Tomato^+^ T cells isolated from colon tumors and non-tumor epithelium and performed single-cell RNA and T cell receptor (TCR) sequencing (scRNA/TCR-seq) using the 10x Genomics Chromium platform to define the transcriptional programs induced by each tumor type. UMAP visualization and unsupervised clustering of fate-mapped CD4^+^ T cells classified them into eight transcriptionally distinct clusters across tumors and colonic IEL (**Figure 4A**). Clusters 0 and 5 corresponded to cytotoxic/ Th1 cells from the IEL and tumor, respectively, marked by the high expression of genes such as *Nkg7, Ccl5, Cxcr3, Tbx21, Ifng, Cd44, Cd226,* and *Bhlhe40*. In contrast, cluster 1 was marked by the expression of *Sell, S1pr1, Igfbp4, Il7r, and Ccr7,* which are classical markers of naïve T cells. Cluster 3 also expressed markers associated with stemness, such as *Tcf7* and *Slamf6.* Clusters 1 and 3 showed downregulation of activation markers, such as *Cd44, Il2ra,* and *Pdcd1* (**Figures 4A–C**). AKP CD4^+^ tumor-infiltrating lymphocytes (TILs) and adjacent IEL were enriched in cytotoxic/Th1 clusters, whereas pro-Met CD4^+^ TILs were predominantly represented in naïve-like clusters 1 and 3 (**Figures 4A–D**). Of note, although our sorting was enriched for ex-naïve CD62L^−^ Tomato^+^ T cells, cells in clusters 1 and 3 retained a naïve-like profile at the transcriptional level.

**Figure 4:**
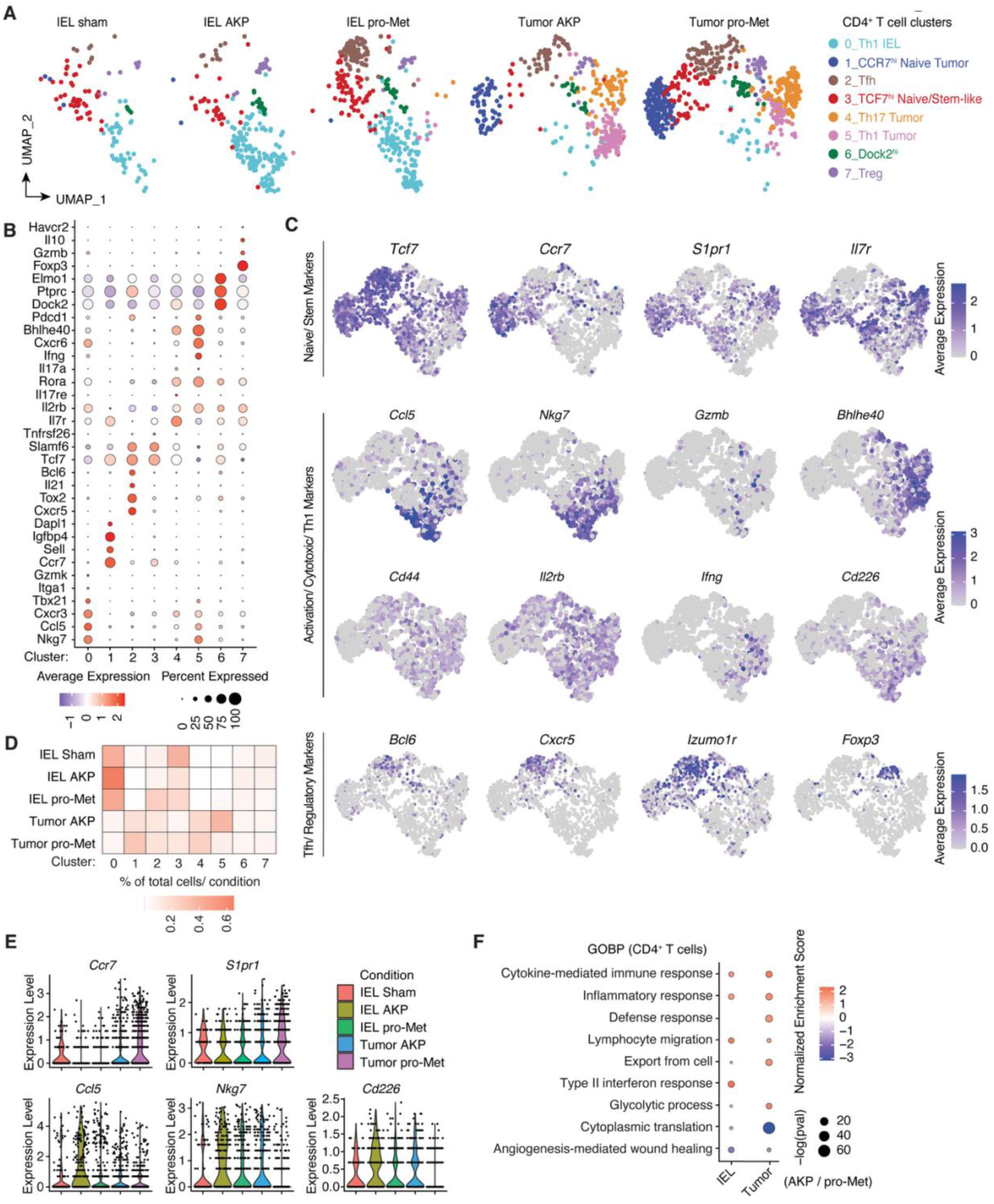
Transcriptional profiles of recently recruited CD4^+^ T cells in early CRC stages. i*Sell*^Tomato^ mice were orthotopically injected with sham, AKP, or pro-Met tumors using a colonoscope 1 day after the second tamoxifen dose. CD62L^−^ Tomato+ TCRαβ^+^ T cells from colon IEL and tumors were subjected to single-cell RNA sequencing. (A) UMAPs of CD62L^−^ Tomato^+^ CD4^+^ T cells across tissues and conditions. Clustering was performed using Seurat (SCT normalization, resolution = 0.6). (B) Dot plot of CD4^+^ T cell clusters visualized in (A), showing the selected markers expressed per cluster. (C) Expression of key marker genes for naïve/stem clusters (*Tcf7, Ccr7, S1pr1, Il7r*), activation/cytotoxic/Th1 clusters (*Ccl5, Nkg7, Gzmb, Bhlhe40, Cd44, Il2rb, Ifng, Cd226*), and Tfh/regulatory clusters (*Bcl6, Cxcr5, Izumo1r, Foxp3*). (D) Heat map showing the frequency of cells per condition across clusters. Each condition (row) represents 100%. (E) Violin plots of selected genes differentially expressed under AKP and pro-Met conditions. Each dot inside the violins represents a cell. (F) Selected Gene Ontology Biological Processes (GOBP) differentially expressed between AKP, pro-Met IEL, and tumor-derived CD4^+^ T cells shown as a dot plot. Colors represent the Normalized Enrichment Score (NES) associated with that pathway in IEL and Tumor and the dot radius represents - log of the adjusted p-value of that enrichment. Single-cell RNA-seq data in all figures are pooled from n=3 mice per condition.

Pro-Met CD4^+^ IEL and TILs were also enriched in cluster 2, which displayed a T follicular helper (Tfh) signature characterized by the expression of *Cxcr5, Bcl6, Il21,* and *Tox2*. T helper 17 (Th17) signature was assigned to cluster 4, marked by *Rorc, Il17a*, and *Il17re,* and was dominated by pro-Met CD4^+^ TILs. Cluster 7 included regulatory T cells (Tregs) defined by the expression of *Foxp3, Il10*, and *Gzmb*. Pro-Met CD4^+^ TILs were slightly more enriched in this cluster than in the AKP CD4^+^ TILs (**Figures 4A–D**).

As indicated by the cluster distributions and selected differentially expressed genes, AKP tumors and adjacent IEL were enriched for activated cytotoxic and Th1 cells, whereas pro-Met tumor CD4^+^ T cells and IEL expressed naïve-like, Tfh, and Th17 signature genes (**Figures 4D, E, and S3A**). Gene Ontology Biological Process (GOBP) enrichment analysis further supported this dichotomy, showing the upregulation of lymphocyte migration, activation, and cytokine-mediated immune response pathways in AKP CD4^+^ T cells, in contrast to the upregulation of cytoplasmic translation and angiogenesis-mediated wound healing in pro-Met CD4^+^ T cells (**Figure 4F**). These analyses suggest that AKP tumors induce robust CD4^+^ T cell activation and cytotoxic differentiation programs that pro-Met tumors fail to induce.

Fate-mapped CD8αβ^+^ T cells were classified into four main clusters. Similar to pro-Met CD4^+^ TILs, pro-Met CD8αβ^+^ TILs displayed a naïve/stem cell-like phenotype. A total of 78% of pro-Met CD8αβ^+^ TILs (compared to 54% of AKP counterparts) localized to cluster 1, characterized by high expression of *Ccr7, Sell, Lef1, S1pr1, Tcf7,* and *Il7r* (**Figures S3B and C).** In contrast, 23% of AKP CD8αβ^+^ TILs (compared to 9% of pro-Met counterparts) localized to a cytotoxic effector cluster 2, marked by the expression of both activation and TCR stimulation markers such as *Ccl5, Nkg7, Nr4a1, Dusp5, Icos, Ctla4, Tigit, Il2rb,* and *Pdcd1* (**Figures S3B and C**), again reflecting a more activated state in AKP tumors. This was further supported by GOBP analysis, which showed the upregulation of T cell activation and inflammatory pathways in AKP tumors (**Figure S3D**). Compared to AKP IEL, pro-Met IEL displayed a cytotoxic profile, as indicated by the expression of genes such as *Gzma, Gzmb, Ifitm1, Ifitm2,* and *Cd226*. Paradoxically, this was accompanied by lower expression of *Ifng*, *Nkg7*, and *Il2rb* (**Figure S3E**). Accordingly, GOBP analysis showed upregulation of granzyme-mediated programmed cell death and signaling and defense response, but also less T cell activation signaling in pro-Met IEL CD8αβ^+^ T cells (**Figure S3D**). These data indicate a lack of CD8αβ^+^ T cell activation in pro-Met tumors.

### Distinct early T cell states in AKP and pro-Met tumors in fate-mapped T cells

scRNA-seq revealed distinct transcriptional profiles discriminating cytotoxic and naïve T cell states in AKP and pro-Met tumors, respectively. Next, we compared both early- and late-stage T cell recruitment phenotypes with pre-existing gut-resident immune responses following the same experimental design shown in Figure 3A. Non-fate-mapped Tomato⁻ CD4⁺ TILs had comparable frequencies of CD62L^+^ cells across tumor models and time points. In contrast, fate-mapped Tomato⁺ CD4⁺ TILs displayed a significantly higher proportion of CD62L^+^ cells in pro-Met than in AKP tumors during the early stages (**Figure 5A**). Both Tomato^−^ and Tomato^+^ CD8αβ⁺ TILs showed stronger enrichment for CD62L^+^ cells in pro-Met tumors than in AKP tumors. However, in the late stage, the proportion of CD62L-expressing cells was similarly high among Tomato⁺ CD4⁺ or CD8αβ⁺ T cells in both tumor models (**Figures 5A and 5B**). Together, these data indicate that the colon contains a baseline pool of CD62L⁺ CD4⁺ and CD8αβ⁺ T cells, likely residing within colonic patches and isolated lymphoid follicles. Fate-mapping these naïve cells residing in the colon and draining lymph nodes during early CRC further showed that these cells differentiated into activated effector states only in AKP tumors, whereas in pro-Met tumors, they largely remained in a naïve state.

**Figure 5:**
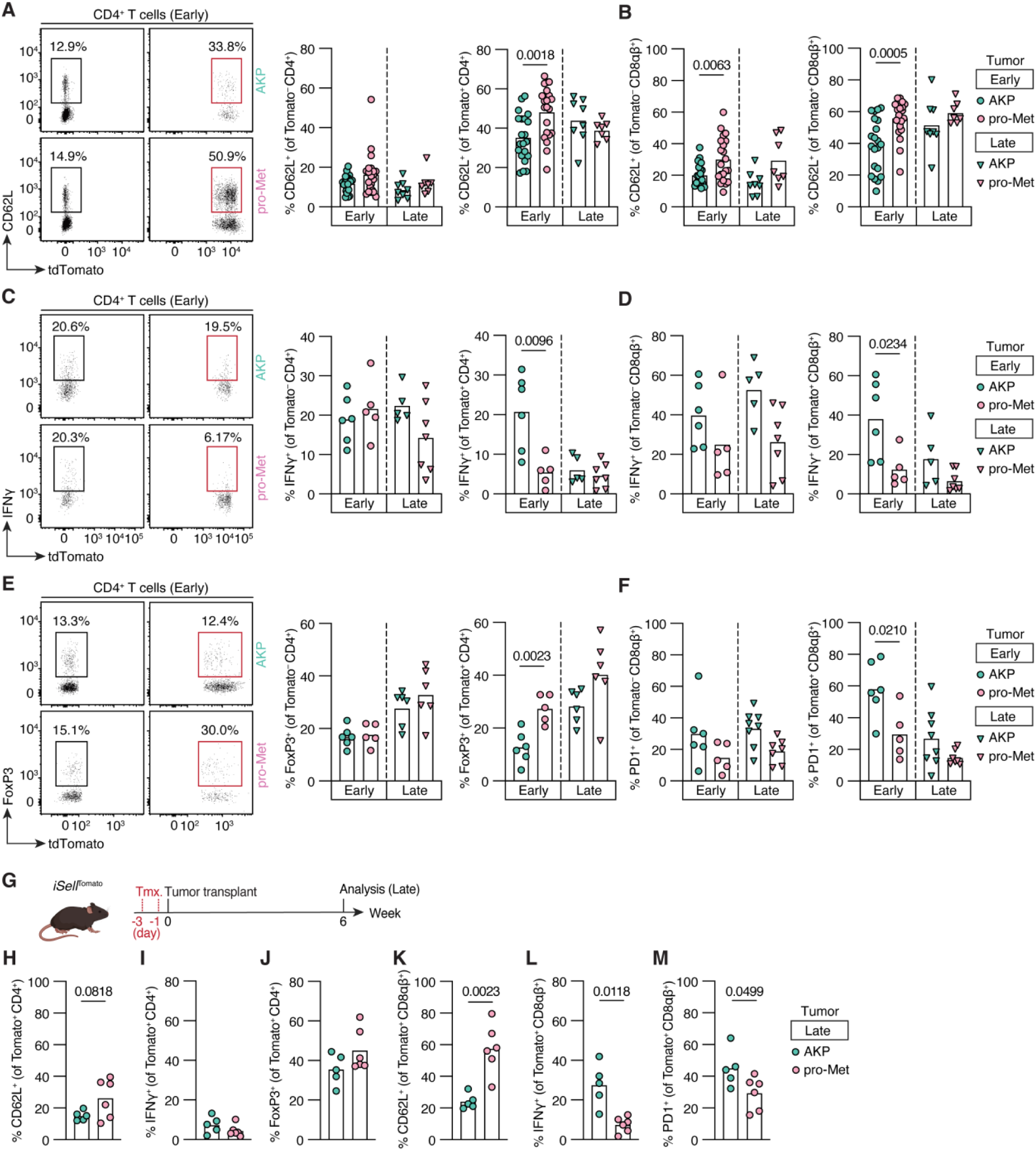
Compromised immune profile of recently-recruited CD4^+^ and CD8αβ^+^ T cells in pro-Met colon tumors. (A-F) Experimental design as shown in Figure 1I: i*Sell*^Tomato^ mice were orthotopically injected with AKP or pro-Met tumors using a colonoscope. Tamoxifen was administered either before tumor injection (early time point) or 4 weeks after tumor injection (late time point). Flow cytometry analysis of Tomato^+^ and Tomato^−^ CD4^+^ and CD8αβ^+^ T cells isolated from colon tumors was performed. (A) Dot plots and frequencies of CD62L^+^ among Tomato^−^ (left) and Tomato^+^ (right) CD4^+^ T cells. (B) Frequencies of CD62L^+^ among Tomato^−^ (left) and Tomato^+^ (right) CD8αβ^+^ T cells. (C) Dot plots and frequencies of IFN-γ^+^ among Tomato^−^ (left) and Tomato^+^ (right) CD4^+^ T cells following *ex vivo* PMA/ionomycin stimulation. (D) Frequencies of IFN-γ^+^ among Tomato^−^ (left) and Tomato^+^ (right) CD8αβ^+^ T cells following *ex vivo* PMA/ionomycin stimulation. (E) Dot plots and frequencies of FoxP3^+^ among Tomato^−^ (left) and Tomato^+^ (right) CD62L^−^ CD4^+^ T cells. (F) Frequencies of PD-1^+^ among Tomato^−^ (left) and Tomato^+^ (right) CD62L^−^ CD8αβ^+^ T cells. (G) Experimental design for H-M. i*Sell*^Tomato^ mice were injected with two tamoxifen doses and orthotopically injected with AKP or pro-Met tumor organoids. Flow cytometry analysis of Tomato^+^ CD4^+^ and CD8αβ^+^ T cells in colon tumors was performed 6 weeks later. (H) Frequencies of CD62L^+^ among Tomato^+^ CD4^+^ T cells. (I) Frequencies of IFN-γ^+^ among Tomato^+^ CD4^+^ T cells. (J) Frequencies of FoxP3^+^ among Tomato^+^ CD62L^−^ CD4^+^ T cells. (K) Frequencies of CD62L^+^ among Tomato^+^ CD8αβ^+^ T cells. (L) Frequencies of IFN-γ^+^ among Tomato^+^ CD8αβ^+^ T cells. (M) Frequencies of PD-1^+^ among Tomato^+^ CD62L^−^ CD8αβ^+^ T cells. Unpaired Student’s t-test was performed between early AKP and early pro-Met, and p-values are indicated. Each symbol represents a mouse.

While IFN-γ secretion among non-fate-mapped TILs (Tomato^−^ CD4^+^ and CD8αβ^+^) was not significantly affected by tumor type or stage, fate-mapped tumor-infiltrating T cells showed higher IFN-γ production in early AKP tumors than in early pro-Met or late-stage tumors (**Figures 5C and D**). Accordingly, FoxP3^+^ Tregs were more frequent among Tomato^+^ CD4^+^ T cells infiltrating early-stage pro-Met tumors than in AKP tumors, whereas no difference was observed in the non-fate-mapped compartment (**Figure 5E**). This early enrichment of FoxP3^+^ Tregs among fate-mapped T cells in pro-Met tumors suggests that the immunosuppressive environment is established at the earliest stages of pro-metastatic tumor development, potentially contributing to the failure of effector T cell differentiation in this model. Additionally, consistent with the reduced activation environment, a smaller proportion of Tomato^+^ CD8αβ^+^ T cells infiltrating pro-Met tumors expressed PD-1 than those in AKP tumors at an early stage, a difference not observed among Tomato^−^ CD8αβ^+^ T cells (**Figure 5F**). Collectively, these findings reinforce that fate-mapped CD4⁺ and CD8αβ⁺ T cells in pro-metastatic CRC acquire an early hypoactivated state, similar to that of newly fate-mapped T cells at the late stages of both AKP and pro-Met CRC models.

To assess how tumor progression influenced the activation state of early recruited T cells, tamoxifen was administered before tumor injection, and fate-mapped Tomato⁺ T cells were analyzed six weeks later (**Figures 5G–M**). CD4^+^ T cells recruited to both AKP and pro-Met tumors showed downregulation of CD62L compared to their levels at the early time point, although CD4^+^ T cells in pro-Met tumors retained slightly higher CD62L expression (**Figures 5A and H**). Moreover, these early fate-mapped CD4^+^ T cells lost their capacity to produce IFN-γ in the AKP model (*p*=0.019), rendering them functionally similar to those in pro-Met tumors (**Figures 5C and I**). Consistently, the frequencies of fate-mapped CD4^+^ Tregs in AKP tumors were comparable to those observed in pro-Met tumors (**Figure 5J**). Early fate-mapped CD8αβ^+^ T cells in AKP tumors maintained a higher activation state at the late time point, as indicated by persistent CD62L downregulation, sustained IFN-γ production, and elevated PD-1 expression. In contrast, CD8αβ^+^ T cells in pro-Met tumors remained in an inactivated state (**Figures 5B, D, F, and K–M**). These data indicate that although AKP tumors induce robust early CD4^+^ and CD8αβ^+^ T cell activation, only CD8αβ^+^ T cells can sustain this early activation state as tumors progress, which may underlie the enhanced primary tumor growth observed upon CD8⁺ T cell depletion.

### AKP tumors induce CD4^+^ T cell clonal expansion

Given the distinct early T cell trajectories observed in AKP versus pro-Met models, we next asked whether these trajectories were associated with divergent clonal composition of tumor-infiltrating CD4⁺ and CD8αβ⁺ T cells. We characterized the CD4⁺ and CD8αβ⁺ T cell repertoire in sham, AKP, and pro-Met models using the scRNA/TCR-seq dataset. Clonal expansion was prominent among CD4⁺ T cells that infiltrated AKP tumors and adjacent non-tumor epithelium (**Figures 6A and S4A**). Accordingly, the clonal overlap was high between the IEL and tumor compartments, with an average MHI of 0.57 (**Figure 6B**). In contrast, pro-Met tumors and IEL CD4⁺ T cells displayed a diverse TCR repertoire, similar to the diversity observed in sham IEL, and thus exhibited less clonal overlap between the two compartments (MHI=0.13) (**Figures 6A, B and S4A**). Cluster distribution analysis revealed that 42% and 40% of the cells in the top 20 expanded CD4⁺ clones were localized to cluster 0 (Th1 IEL) and cluster 5 (Th1 Tumor), respectively (**Figures 6C and D**). The remaining CD4⁺ T cells in the top 20 expanded clones were distributed among Th17 cluster 4 (8%), Dock2^hi^ cluster 6 (5%), Tfh cluster 2 (2%), and TCF7^hi^ naïve/stem cluster 3 (2%). This distribution aligned well with the transcriptional profiles of these clusters, with most expansions occurring in the cytotoxic and activated clusters. Additionally, 16 of the top 20 expanded clones were from AKP tumors and IEL, whereas the remaining four were from pro-Met IEL and tumor-infiltrating T cells (**Figure 6D**). These data confirm that non-metastatic AKP tumors are more immunogenic than pro-Met tumors, leading to robust CD4⁺ T cell clonal expansion.

**Figure 6:**
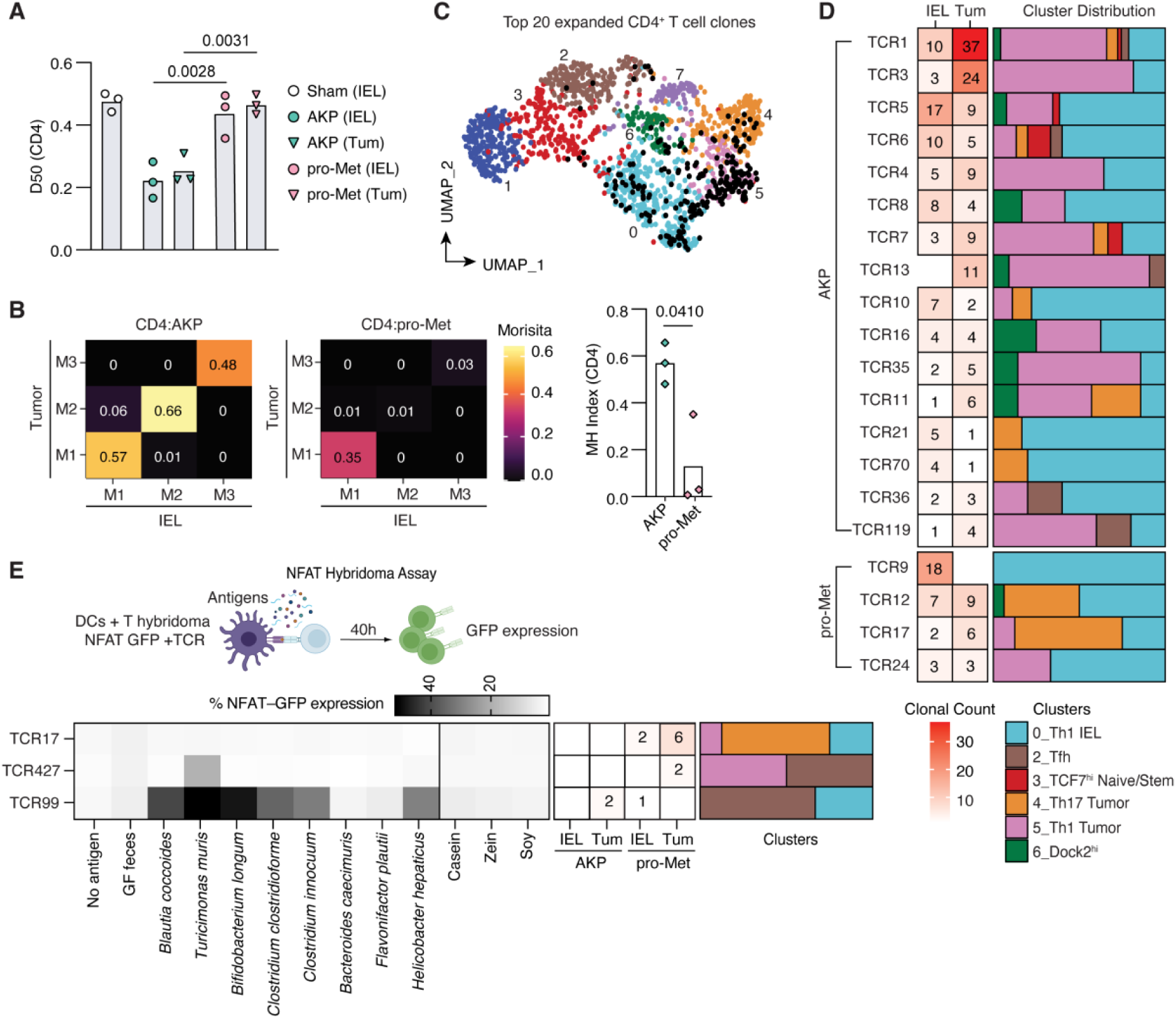
AKP tumors induce clonal expansion of CD4^+^ T cells in colon tumors and epithelium. i*Sell*^Tomato^ mice were orthotopically injected with sham, AKP, or pro-Met tumors using colonoscope 1 day post second tamoxifen dose. CD62L^−^ Tomato+ TCRαβ^+^ T cells from colon IEL and tumors were isolated for single-cell RNA and TCR sequencing two weeks after orthotopic injections (early time point). (A) Diversity index 50 (D50) of CD4^+^ T cell clones isolated from indicated tissues and conditions. Each symbol represents a mouse. (B) CD4^+^ T cell clonal sharing between the IEL and tumor in AKP (left) and pro-Met (right) injected mice, represented by the Morisita-Horn (MH) index. M1,2,3 = Mouse 1,2,3. Each symbol represents a mouse. (C) Top 20 expanded CD4^+^ T cell clones (black) overlayed on CD4 UMAP. Seurat-identified clusters (resolution =0.6) were visualized and labelled on the UMAP. (D) Clone and cluster distribution of top 20 expanded CD4^+^ T cell clones isolated from AKP or pro-met tumors. The top 16 clones were expanded in AKP-injected mice, and the last four clones were found in pro-Met-injected mice. Numbers inside squares represent the number of cells detected for each clone in the IEL or tumor (Tum). Left, the cluster distribution of each clone. (E) TCRs from expanded CD4^+^ T cell clones were reconstructed in TCR-deficient murine NFAT-GFP T cell hybridomas and co-cultured with antigen-pulsed splenic dendritic cells (DCs) in the presence of the indicated antigens (bacterium species or dietary proteins) for 40 h. Frequency of NFAT-GFP expression indicating TCR reactivity is represented on the heatmap. Numbers inside colored squares represent the number of cells detected for each clone in the IEL or tumor (Tum), with cluster distribution corresponding to each clone. Ordinary one-way ANOVA with Tukey’s multiple comparison test was performed in (A), and unpaired Student’s t-test in (B).

CD8αβ⁺ T cells exhibited distinct clonal dynamics compared to CD4⁺ T cells. First, no clonal expansion was observed in tumor-infiltrating CD8αβ⁺ T cells in the AKP or pro-Met models. However, moderate clonal expansion was observed in IEL CD8αβ⁺ T cells in both the AKP and pro-Met groups; their D50 was only slightly lower than that observed in the sham control (**Figure S4B**). Additionally, there was minimal clonal overlap between the tumor and IEL compartments in both tumor models (**Figure S4C**). The majority (90%) of the top 20 expanded CD8αβ⁺ T cell clones were assigned to the cytotoxic IEL cluster 0 (**Figure S4D**).

Given the pronounced clonal expansion of CD4⁺ T cells observed in mice injected with AKP organoids, we next sought to identify the TCR specificities of these expanded clones. We selected 21 CD4⁺ T cell clones, reconstructed their TCRs in NFAT–GFP hybridomas, and confirmed their functional capacity to respond to TCR stimulation following anti-CD3 treatment (**Figures S4E, F and Supplementary Table 1**). We then tested the hybridomas that showed robust anti-CD3 responses for reactivity against tumor, microbial, and dietary antigens using an *in vitro* screening assay^37,38^ (**Figure 6E**). Whereas none of the clones tested recognized tumor antigens, one clone (TCR17) showed reactivity to a standard chow diet, and another clone (TCR99) reacted to feces collected from SPF control, AKP, or pro-Met tumor-bearing mice (**Figure S4F**). To narrow down the dietary and microbial candidates, we screened these clones for reactivity against defined dietary proteins and commensal species present in the Oligo-MM^12^ microbiota, which encompasses major species found in the SPF microbiota^39^, including those in our facility (**Figure 6E**). We also tested reactivity against *Helicobacter hepaticus,* which has been previously associated with antitumor immunity in a CRC model^40^. The dietary-reactive Clone TCR17, which was detected in pro-Met tumors and IEL, did not show reactivity to the purified casein, zein, or soy proteins (**Figure 6E**). Among the TCR clones tested for specific microbes, TCR427, which was detected in pro-Met tumors, responded only to *Turicimonas muris* lysates. In contrast, TCR99 recognized various bacterial species, including *Blautia coccoides, Turicimonas muris, Bifidobacterium longum, Clostridium clostridioforme, Clostridium innocuum,* and *Helicobacter hepaticus* (**Figure 6E**). However, TCR99 did not respond to *Flavonifactor plautii* or *Bacteroides caecimuris,* suggesting broad but selective reactivity. TCR99 is a public clone shared between AKP tumors and pro-Met IEL. Notably, the two TCR clones responding to microbial antigens (TCR427 and TCR99) were localized mostly in the Tfh cluster 2, in addition to the Th1 clusters 0 and 5 (**Figure 6E**). Given the microsatellite stability of the organoids utilized, the observed T cell expansion and antitumor activity likely occur independently of classical neoantigen recognition, potentially through cross-reactive T cells or antigen-independent engagement of stress-induced molecules^8,41^. The patterns of TCR reactivity indicate that antimicrobial and dietary responses contribute to the composition of the T cell repertoire during tumor progression. Overall, these results uncover marked T cell expansion in response to AKP tumors that was coupled with effector and antitumor T cell differentiation. T cell clonal expansion was in part directed towards non-tumor antigens derived from diet and microbiota, suggesting a role for the steady-state homeostatic immune repertoire in the tumor responses.

### Tumor-intrinsic MHCII expression modulates tumor outcome

Having defined the clonal architecture and broad antigen specificities of the tumor-induced repertoire, we next addressed whether fate-mapped T cells are functionally capable of controlling tumor cell growth, or whether this capacity differs with metastatic potential. We established a tumor organoid co-culture system using T cells isolated from the tumor and adjacent epithelium of i*Sell*^Tomato^ mice two weeks after orthotopic transplantation with either AKP or pro-Met tumors. Following our *in vivo* fate-mapping strategy (*see* Figure 3A), we sorted CD4^+^ and CD8^+^ T cells into three distinct subsets: naïve or central memory (CD62L^+^), resident effector or memory (CD62L^−^) Tomato^−^ cells, and recently activated, fate-mapped effector (CD62L^−^) Tomato^+^ cells. These populations were then co-cultured with dissociated pro-Met tumor organoids and organoid recovery, cytokine release, and changes in tumor cell phenotype were quantified five days later (**Figure 7A**). Only fate-mapped T cells derived from the non-metastatic AKP tumor environment demonstrated antitumor activity, reducing the number of healthy pro-Met organoids compared to other conditions (**Figures 7B–D, and S5A**). This suppressive effect was specific to the non-metastatic TME, as fate-mapped CD62L^−^ Tomato^+^ T cells derived from pro-Met tumors failed to control organoid growth (**Figures 7B–D, and S5A**).

**Figure 7:**
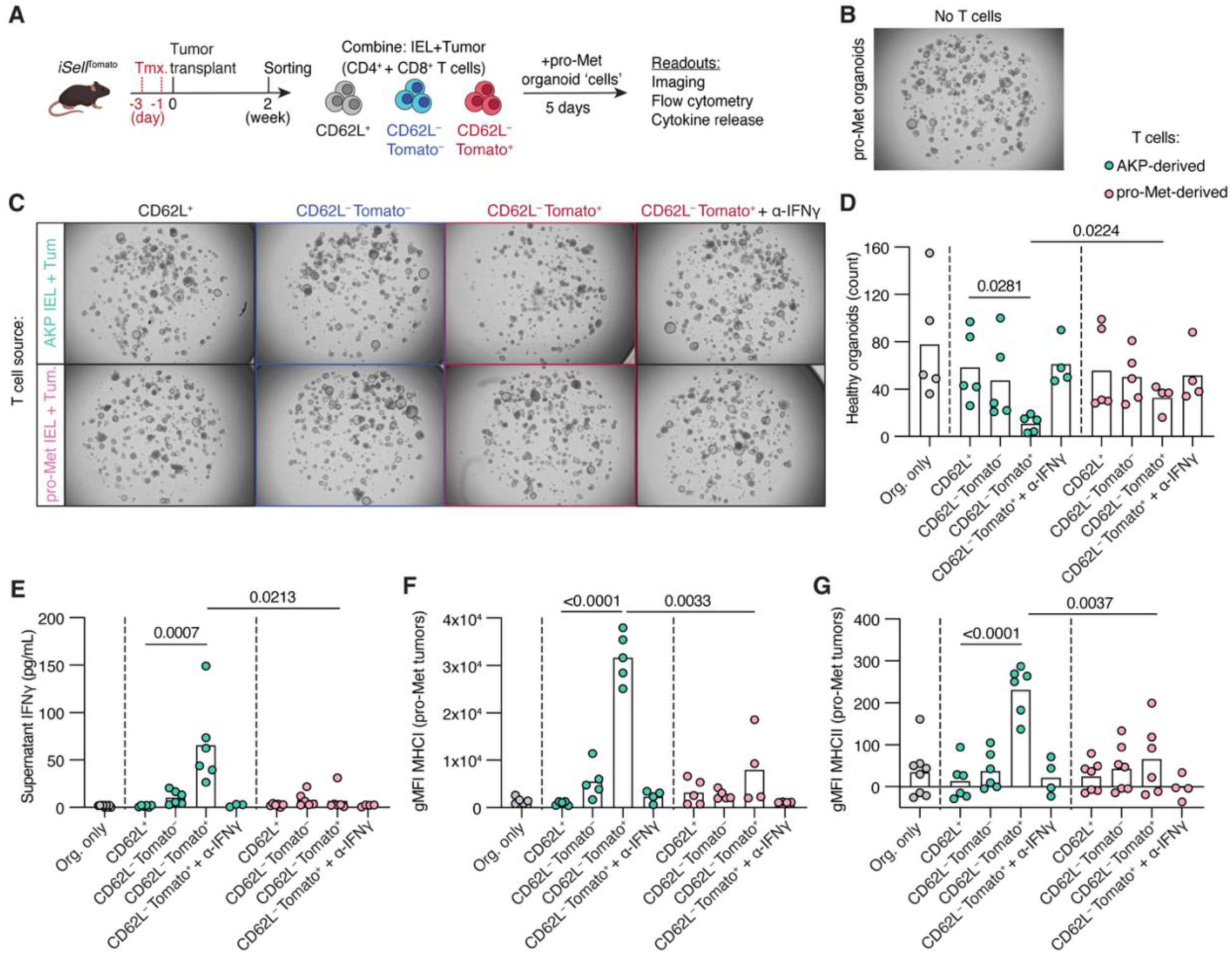
Fate-mapped, AKP-derived TCRαβ^+^ T cells suppress tumor growth *in vitro* in an IFN-γ-dependent manner. (A) Co-culture experimental design. i*Sell*^Tomato^ mice were injected with two doses of tamoxifen on days - 3 and −1, followed by colonoscopy-guided orthotopic transplantation of AKP or pro-Met tumor organoids on day 0. Colon tumors and IEL were harvested two weeks later, and the indicated cell populated were sorted. For each condition, approximately 4500 T cells were co-cultured in matrigel for five days with 1000 pro-Met organoid cells. Data is pooled from 3 independent experiments. (B) BF imaging (2X, 1.5 Zoom) of control pro-Met organoids with no T cell treatment. (C) BF imaging (2X, 1.5 Zoom) of AKP-derived T cell subsets (top) or pro-Met-derived T cell subsets (bottom) pooled from both IEL and tumors of n=4 or 5 mice per condition per experiment. (D) Quantification of healthy organoid numbers per condition. Each dot represents a well. (E) CBA analysis of co-culture supernatant collected on day 5. Graph shows levels of IFN-γ. (F–G) gMFI of MHCI (F) and MHCII (G) expression assessed by flow cytometry on pro-Met tumor organoids harvested at day 5. Ordinary one-way ANOVA with Tukey’s multiple comparison test was performed in (D–G) among AKP-derived and pro-Met-derived T cell conditions. Additionally, unpaired Student’s t-tests were performed between AKP-derived and pro-Met-derived CD62L^−^Tomato^+^ T cell conditions.

To investigate the mechanism driving this antitumor activity, we analyzed the cytokine profile in the co-culture supernatants. We detected secretion of IFN-γ exclusively in the co-cultures containing the cytotoxic, AKP-derived fate-mapped T cells (**Figure 7E**), while no significant differences in TNF-α secretion were observed in any of the conditions (**Figure S5B**). The addition of an anti-IFN-γ blocking antibody abrogated this cytotoxic effect, confirming that the observed tumor cell killing was dependent on IFN-γ (**Figures 7C–E**). IFN-γ is a pleiotropic cytokine known to increase tumor cell visibility by upregulating antigen presentation machinery, including MHCI and MHCII. Consistent with this, we observed an upregulation of MHCI and MHCII surface expression on the organoid cells specifically in the presence of the IFN-γ-producing, AKP-derived fate-mapped T cells (**Figure 7F and G**). These findings functionally validate the i*Sell*^Tomato^ model, demonstrating that it specifically enriches for tumor-induced T cells capable of IFN-γ-mediated tumor suppression, a functional state that is impaired in T cells recruited to pro-metastatic tumors.

The divergent T cell responses induced by AKP and pro-Met tumors prompted us to ask whether this reflects underlying genomic differences between the two lines. To address this, we performed Whole Exome Sequencing (WES) on both organoid lines. Strikingly, the two lines carried a highly comparable mutational burden, sharing 1,102 somatic variants, with only 137 AKP-specific and 129 pro-Met-specific mutations identified, with each representing less than 12% of the total mutational load (**Supplementary table 2**). The expected driver mutations in *Apc, Kras*, and *Tp53* were confirmed in the shared mutation set, consistent with the common clonal origin of both lines. Among the pro-Met-specific mutations, we found several alterations in genes related to immune regulation, including mutations in *Hivep1*, *Ppp2r5a*, *Sp110*, and *Nucks1*, which may influence immune-related gene expression and chromatin-associated regulatory programs (**Supplementary table 2**). Nevertheless, no mutations were detected in core antigen presentation genes, and the overall mutational divergence between the lines remained modest. These data suggest that the functional differences between AKP and pro-Met are unlikely to be driven by gross genomic divergence and instead point toward transcriptional or epigenetic reprogramming as the primary mechanism of immune evasion. Consistent with this, pro-metastatic reprogramming in CRC has been shown to be driven by epigenetic plasticity, including the emergence of fetal-like and basal-like/squamous transcriptional programs converging on GATA6 loss, rather than by additional genetic alterations^42^.

Given the prominent CD4^+^ T cell clonal expansion observed exclusively in the AKP model, suggesting a productive MHCII-dependent antigen presentation, we directly examined MHCII expression on tumor cells from both models. Flow cytometric analysis of early-stage *ex vivo* tumors revealed a tumor-specific reduction in MHCII expression on pro-Met tumor cells compared to AKP tumors, while MHCII expression on the adjacent colon epithelium remained comparable between the two models (**Figure 8A**). To directly test whether this loss of MHCII is a causal driver of immune evasion and metastasis, we generated an MHCII-deficient AKP organoid line by knocking out Class II transactivator (*Ciita*) using CRISPR/Cas9 (AKP.*Ciita*^KO^) (**Figures 8B and S5C**). While early primary colon tumor sizes were comparable between mice transplanted with AKP.*Ciita*^KO^ or its parental wild-type AKP line (**Figure 8C**), the loss of MHCII resulted in a reduction in the frequency of tumor-infiltrating CD4^+^ T cells (**Figure 8D**). Furthermore, scTCR-seq analysis revealed that CD4^+^ T cells infiltrating AKP.*Ciita*^KO^ tumors exhibited a higher D50 compared to those in wild-type AKP tumors, indicating a failure to undergo clonal expansion characteristic of the responses induced by AKP tumors (**Figures 8E and S5D**). At the late stage of tumor progression, mice bearing AKP.*Ciita*^KO^ tumors developed significantly larger primary colon tumors than those bearing wild-type AKP tumors, and approximately 28% of AKP.*Ciita*^KO^ mice developed liver macrometastases. In contrast, none of the mice transplanted with the parental AKP model developed metastasis (**Figure 8F**). In a complementary experiment, overexpressing *Ciita* in pro-Met organoids (pro-Met.*Ciita*^OE^) did not alter late-stage colon tumor size or liver metastasis (**Figure S5E**), suggesting that while MHCII loss is critical, restoring it alone is insufficient to overcome the multifaceted immune evasion programs established in pro-metastatic tumors. Finally, we assessed the impact of MHCII loss on physical tumor–T cell interactions using the uLIPSTIC system. The proportion of tumor-interacting biotin^+^ CD4^+^ or CD8αβ^+^ T cells was unchanged in AKP.*Ciita*^KO^ compared to wild-type AKP tumors, and similarly, pro-Met.*Ciita*^OE^ did not rescue T cell interactions relative to wild-type pro-Met tumors (**Figure S5F**), This suggests that physical tumor–T cell interactions are governed by MHCII-independent mechanisms in both models. Together, these findings identify tumor-intrinsic MHCII as a key regulator of CD4⁺ T cell responses that restrains primary tumor growth.

**Figure 8:**
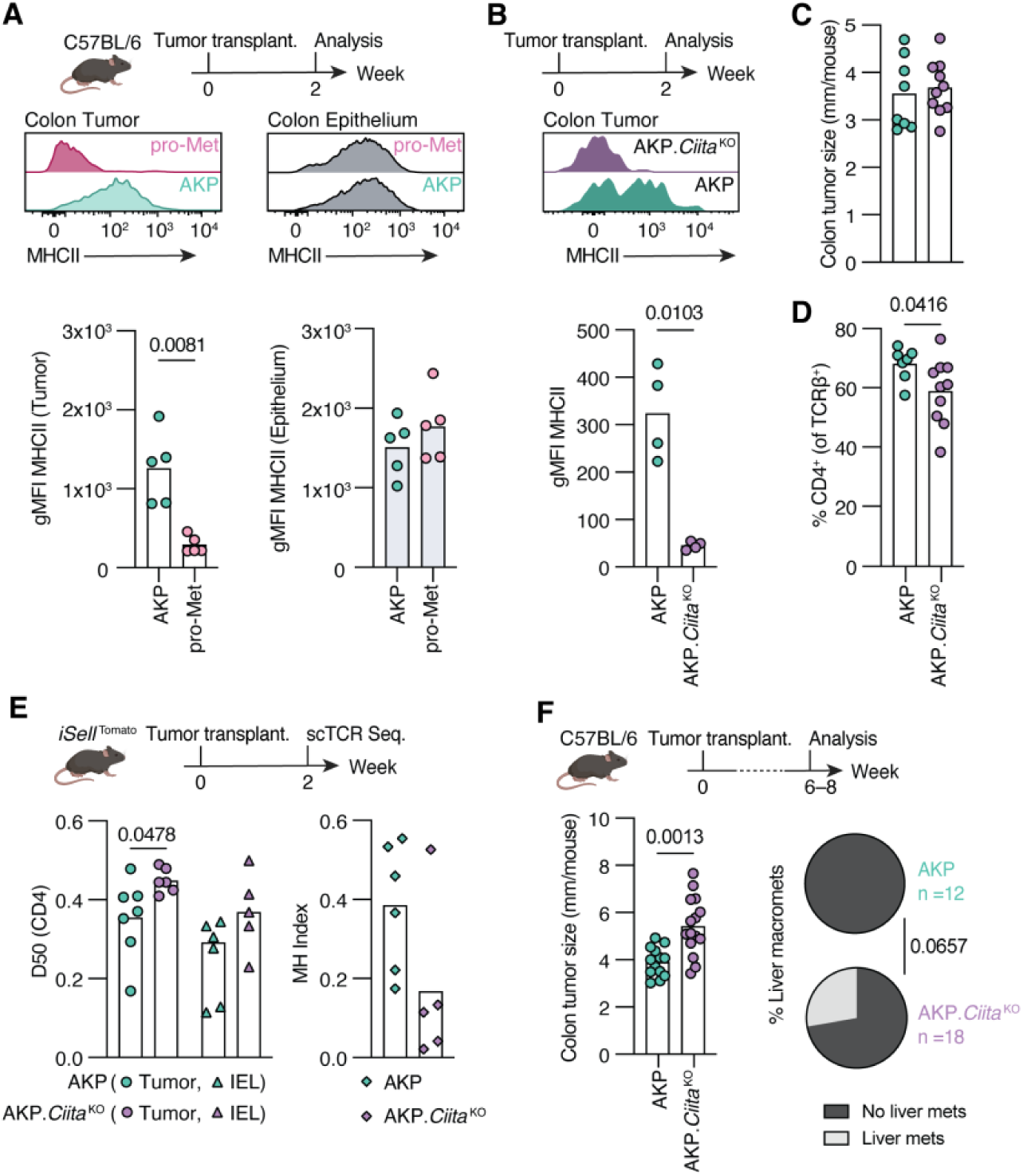
AKP tumor-intrinsic MHCII modulates CD4^+^ T cell repertoire and restricts tumor growth. (A) gMFI of MHCII in AKP and pro-Met *ex vivo* tumors and nearby colon epithelium harvested two weeks post tumor transplantation. (B) Orthotopic transplantation of AKP or AKP.*Ciita* ^KO^ tumor organoids and validation of MHCII downregulation *ex vivo*. (C) Average colon tumor size per mouse two weeks post tumor transplantation. (D) Frequency of CD4^+^ T cells of AKP and AKP.*Ciita* ^KO^ tumors two weeks post transplantation using flow cytometry. (E) scTCR seq of fate-mapped CD62L^−^ Tomato^+^ CD4^+^ T cells harvested from colon tumors and nearby IEL of indicated tumors, and MH index indicating clonal overlap between tumor and IEL compartments. (F) Average colon tumor size per mouse (left graph), and pie charts representing liver macrometastases at the late tumor stage (6-8 weeks post transplantation). AKP control shown in (F) is from the same dataset in Figures 1B and 1C. Ordinary one-way ANOVA with Tukey’s multiple comparison test was performed for colon tumor size with all groups combined. Fisher’s exact test was performed for liver macrometastases data. Unpaired Student’s t-tests were performed in (A–E).

## Discussion

We provided a detailed characterization of CD4^+^ and CD8^+^ T cell clonality, activation states, and cellular interactions across CRC models with distinct metastatic potential. Using orthotopic CRC models coupled with temporal T cell fate mapping, we uncovered divergent early activation trajectories of tumor-infiltrating T cells that reflect the metastatic potential of tumors. Non-metastatic AKP tumors elicited a strong cytotoxic and Th1-skewed immune response in recently recruited T cells, accompanied by enhanced CD4^+^ T cell clonal expansion. This contrasts with the naïve-like and regulatory phenotypes of T cells recruited to pro-metastatic tumors during the same early window, suggesting an accelerated state of immune hypoactivation^13,43^. These findings contribute to the growing understanding that antitumor immunity in CRC is heterogeneous, varying widely with cancer stage, tumor-intrinsic factors, and tumor environmental cues, and is critical in predicting disease outcomes.

Previous studies have shown that the overall immune infiltrate and its spatial organization, defined as Immunoscore, outperform microsatellite instability status in predicting patient survival^44–46^. Our data further demonstrate that both the quality and quantity of early T cell infiltration and activation differ markedly between non-metastatic and pro-metastatic settings. Additionally, early recruited T cells, particularly CD8^+^ T cells, retained their initial activation state as CRC progressed, whereas CD4^+^ T cells exhibited a gradual decline in their activation state. These findings resonate with recent observations in human CRC, where early stage pMMR tumors displayed robust T cell infiltration and activation that diminished as tumors advanced and metastasized^47^. These findings reflect the importance of early immune activation and suggest that it may predict cancer outcomes.

The distinct CD4^+^ and CD8^+^ T cell clonal dynamics explored in our study have multiple important implications. First, the distinct CD4^+^ T cell clonal expansion observed in the two tumor models indicates that differences in TCR clonality may be associated with disease outcome. A previous study reported that increased TCR clonal expansion in metastatic melanoma patients was predictive of their response to ICB treatment and overall survival^48^. Another study suggested that ICB treatment can increase clonal expansion in pMMR CRC patients to levels comparable to dMMR CRC, highlighting that localized clonal expansion may still occur in pMMR tumors^47^. These studies evaluated TCR clonality considering total tumor-infiltrating T lymphocytes without distinguishing between CD4^+^ and CD8^+^ T cell subsets^47,48^. Our data revealed that CD4^+^ T cells undergo greater clonal expansion than CD8^+^ T cells in the non-metastatic AKP model. This points to a potential contribution of certain CD4^+^ T cell clones in shaping antitumor immunity, although the outcomes of CD4^+^ and CD8^+^ T cell depletion or immune-deficient experiments suggest that the repertoire alone cannot determine the metastatic outcomes.

In addition to the TCR clonal dynamics observed in tumor tissues, our findings revealed extensive clonal overlap between CD4^+^ T cells infiltrating tumors and adjacent non-tumor colon epithelium, particularly in the non-metastatic AKP model. These findings, including the microbiota- and food-antigen specificities of some of our expanded clones, suggest that intestinal antitumor responses can co-opt pre-existing TCR repertoires utilizing ongoing mucosal immune circuits to either potentiate or subvert antitumor responses. This hypothesis is further supported by a recent study demonstrating that colonization with segmented filamentous bacteria (SFB) enhances ICB efficacy in mice bearing B16 tumors expressing SFB-derived antigens^49^. This enhancement was mediated by the reprogramming of SFB-specific “ex-Th17” CD4^+^ T cells toward a Th1-like phenotype characterized by IFN-γ and TNF-α secretion, which in turn boosted CD8^+^ T cell cytotoxicity^50^. Another study reported that colonization with *Helicobacter hepaticus* reduced tumor burden in an AOM/DSS CRC model by inducing peritumoral tertiary lymphoid structures that supported tumor regression^40^. These studies further support a previous report that human lung and colorectal tumors are abundantly infiltrated by non-tumor-specific, so-called bystander CD8^+^ T cells^41^. Collectively, these data expand our understanding of antitumor immune responses and point to novel therapeutic avenues.

Although multiple immune evasion mechanisms have been described^51^, the lack of direct tumor–T cell interaction implies potential defects in immune synapse formation, antigen processing and presentation pathways, or alterations in cell ligands^51^. For instance, tumor cells may acquire genetic or epigenetic defects that lead to downregulation of surface MHCI and MHCII expression, impairing T cell recognition^52^. Our finding that tumor-intrinsic MHCII expression modulates the early CD4⁺ T cell response suggests a similar mechanistic layer, placing antigen presentation by the tumor cell itself, and not solely by professional antigen-presenting cells, as a determinant of productive CD4⁺ T cell engagement in CRC ^53–55^. Nevertheless, restoring MHCII in pro-metastatic organoids was not sufficient to rescue T cell interactions or alter tumor outcome, indicating that MHCII downregulation operates within a broader immune-evasion program rather than as a single decisive switch. Tumor MHCII expression has been linked to CD4⁺ T cell–dependent antitumor responses and improved outcomes across several human cancers ^53,56,57^, and our results suggest that its early loss may be one of the initiating events distinguishing metastasis-prone CRC.

Besides disruption to antigen presentation, cancer cells can impact granzyme and perforin-mediated cytotoxic killing through various molecular adaptations, such as increasing cell softness and remodeling actin cytoskeleton to physically impair proper immune synapse formation^58,59^. These studies suggest that altering the metastatic fate of a tumor type likely requires the reversal of multiple evasion mechanisms. For example, we showed that depletion of CD4^+^ or CD8^+^ T cells alone did not induce metastatic behavior in the AKP tumor line, indicating that additional tumor-intrinsic mutations or microenvironmental alterations beyond immune modulation are likely required to confer a metastatic phenotype. These findings also add to the growing appreciation of the role of non-tumor-specific T cells in potentiating tumor outcomes. Early clinical studies dating back to 1976 showed that intravesical administration of Bacillus Calmette–Guérin (BCG), an attenuated *Mycobacterium bovis* vaccine originally developed for tuberculosis, markedly reduces recurrence and progression of non–muscle-invasive bladder cancer, establishing it as the first successful immunotherapy for a solid tumor^60,61^. More recent studies have reported that COVID-19 mRNA vaccination can lead to spontaneous tumor regression by activating type I IFN program, resulting in improved survival in patients with non–small cell lung cancer or metastatic

melanoma^62–64^. Elucidating the mechanisms that govern the immune landscape and mediate tumor-T cell interactions could reveal new therapeutic targets and inform strategies to enhance antitumor immunity. Together, these results define a temporal process in which the quality of the early T cell response distinguishes CRC with and without metastatic potential. While this early immune divergence is not by itself sufficient to dictate metastatic fate, modulation of tumor-intrinsic MHCII represents one node coupling CD4⁺ T cell engagement to the control of primary tumor growth which could influence metastatic dissemination. These findings suggest that the failure of productive local immune engagement is an early marker of metastasis-prone CRC, and that restoring early T cell immunity, enhancing tumor immunogenicity, or leveraging the mucosal antigenic environment may offer new therapeutic strategies.

### Limitations of the study

Our study was based on orthotopic models that reproduce key features of primary and metastatic CRC, although they do not capture the earliest stages of tumorigenesis. Furthermore, the *in vivo* passaging process used to derive the pro-Met line from the AKP line likely selected mutations that increased metastatic capacity while concurrently overriding antitumor immunity in the pro-Met line. Although additional CRC models, such as MC38 and MC38 LVM2, were examined, differences in aggressiveness, kinetics, and metastatic behavior limited direct comparisons. Additionally, our fate-mapping approach effectively identified approximately 60% of recently recruited T cells; hence, incomplete labeling may underreport rare activation patterns. We also acknowledge that the CD62L^−^ Tomato^+^ population is heterogeneous and may include ex-naïve T cells that were activated outside the TME before subsequently migrating into the tumor. Therefore, not all labeled cells can be assumed to have been primed in response to tumor-derived antigens. Moreover, the identity of the antigens recognized by several top expanded clones remains unknown, precluding a definitive link between the observed clonal expansion and antitumor immunity.

## Supporting information

Resource Table

## Author contributions

D.M. conceived the study. M.S., A.M.B, and D.M. initiated the study, designed experiments, and wrote the manuscript. M.S. and A.M.B. performed and analyzed experiments. I.M. and A.R. performed bioinformatics analyses and assisted with interpretation of sequencing data. A.T. performed NFAT hybridoma co-culture assays. Y.S.Y. performed IF staining and imaging. Y.S.Y. and R.C. assisted with flow cytometry experiments. S.E.C. performed histopathological evaluation and interpretation. P.W.D. assisted with uLIPSTIC experiments. N.G. generated and provided the AKP and pro-Met tumor organoids. S.F.T. provided the MC38 cell lines and guided experiments. A.M.B and D.M. supervised the research. All authors edited the manuscript.

## Acknowledgements

We are grateful to Ö. Yilmaz and N. Goto for generating and providing the AKP and pro-Met tumor organoid lines. We thank J. Moltke for providing the *Pou2f3*^CreERT2-IRES-EGFP^ mice, G. Victora for providing the sortase A plasmid constructs, and S. Grivennikov for guidance on tumor organoid models and culture methods. We thank S. Beyaz for providing CIITA virus vector and for valuable feedback. We thank Y. Alvares for assistance with colonoscopy setup and training, and G. L. Reis for help with experiments. We thank A. Rogoz, G. Scrivanti, L. Bianco, Y. Khan, the Comparative Bioscience Center, the genomics and imaging cores, and additional Rockefeller University staff for continuous support. We thank RU Bioinformatics core for helping with WES data analysis. We are grateful to K. Gordon and J.Truman for assistance with cell sorting, and to all members of the Mucida and Victora laboratories for insightful discussions. We thank A. Kamphorst and G. Donaldson for critical review of the manuscript and helpful suggestions. We thank the Laboratory of Comparative Pathology for histopathology support (with funding from NIH Core Grant **P30 CA008748-59**). This work was supported by NIH grants **R01DK093674, R01DK113375**, and **U54CA261701**, the CZI Science, Weill Cancer Hub East, and The Howard Hughes Medical Institute (D.M.).

## Notice of pre-existing conditions, requirements, and licenses for article submission

The article submitted together with this notice is subject to the Immediate Access to Research policy of the Howard Hughes Medical Institute (“HHMI”). In accordance with this policy: (i) a preprint of this article either has been, or will be, deposited on a preprint server under a Creative Commons Attribution 4.0 International (CC BY 4.0) license and (ii) an additional author-published revised version of this article incorporating peer review feedback and/or new results or analysis either has been, or prior to journal publication will be, deposited on a preprint server under a CC BY 4.0 license. In addition, a non-exclusive CC BY 4.0 license to this article has been granted to the public and HHMI has a sublicensable, non-exclusive license to this article. THIS ARTICLE IS SUBMITTED FOR REVIEW AND ACCEPTANCE SUBJECT TO THESE PRE-EXISTING CONDITIONS, REQUIREMENTS AND LICENSES. If you have any concerns with any of these pre-existing conditions, requirements or licenses, please contact the corresponding author immediately

## Data Availability Statement

Raw fastq for the single cell RNA-seq is available at Sequence Read Archive. Processed scRNA-seq data is available on GEO (GSE311111). Raw data for scTCR-seq and WES will be available before publication.

## Methods

### Mice

Animal care and experimentation were consistent with NIH guidelines and were approved by the Institutional Animal Care and Use Committee at The Rockefeller University (IACUC Protocol 25036-H). C57BL/6J (000664), *Rosa26*^CAG–LSL-tdTomato-WPRE^ (007914), *Rag1^−/–^* (002216) mice were purchased from Jackson Laboratories and housed in our facility. *Sell*^CreERT2^ mice were provided by Michel Nussenzweig (Rockefeller)^32,33^. *Pou2f3*^CreERT2-IRES-EGFP^ (037511) mice were provided by Jakob von Moltke (U of Washington)^65^. *Rosa26^uLIPSTIC^* (038221) mice were provided by Gabriel Victora (Rockefeller)^26^. Several of these strains were interbred in our facility to obtain the final strains described elsewhere in the text. All strains were generated and maintained on C57BL/6J background, and orthotopic transplantation experiments with C57BL/6 tumor lines were performed on syngeneic recipient mice. Mice were housed in Specific Pathogen Free conditions on a fixed 12-hour light/dark cycle with food and water provided *ad libitum*. Adult male and female mice (8-16 weeks old) were used in the experiments unless otherwise specified.

### Tumor organoids and cell lines

Naïve AKP organoids^24^ (referred to here as AKP) and pro-Met organoids were generated and provided by Ömer Yilmaz (MIT). Pro-Met organoids were generated by three *in vivo* passages of naïve AKP organoids. Specifically, naïve AKP organoids were orthotopically injected in the colon and animals were kept until liver metastases were detected. Liver tumors were isolated, digested, and expanded *in vitro* as organoids as previously described for organoid generation^24^. This process was repeated twice to generate the tertiary metastatic AKP organoids (referred to here as pro-Met). After each *in vivo* passage, organoids were generated from liver tumors 6-10 weeks after orthotopic injection.

All tumor organoids were cultured by resuspending organoid cells in 60% matrigel in cold ADF^+^ media (Advanced DMEM F12 supplemented with 5% heat inactivated fetal bovine serum (FBS), 1% Penicillin-Streptomycin-Glutamine, and 1% HEPES). In a 6-well plate, 40 uL droplets of cell suspension were plated and incubated at 37°C for 20 min, followed by adding 3 mL of pre-warmed ADF^+^ media.

MC38 and MC38 LVM2 were provided by Sohail Tavazoie (Rockefeller). MC38 LVM2 was generated by passaging MC38 *in vivo* twice: MC38 was intrasplenically injected in mice, and individual liver nodules were dissected after growth, cut into small pieces, and digested in 1 mg/mL Collagenase Type VIII in serum-free DMEM for 30-45 mins while shaking at 37°C. Single cells were filtered through a 70 μm cell strainer into single cell suspension. The cells were resuspended in 5 mL ACK lysis buffer and left at RT for 5 mins. After a PBS wash, the cells were expanded *in vitro* before reinjection into mouse spleen. Secondary liver tumors were isolated and expanded, making the MC38 Liver Metastasis 2 (LVM2). MC38 and MC38 LVM2 cell lines were cultured in DMEM^+^ media (DMEM supplemented with 10% heat inactivated FBS, 2mM glutamine, 0.1 mM nonessential amino acids, 1 mM sodium pyruvate, 10 mM HEPES, 50ug/ml gentamycin sulfate, and 1% Penicillin-Streptomycin).

### Generation of SrtA-organoids

To produce SrtA-expressing retrovirus, 4×10^5^ BOSC23 cells^66^ resuspended in 2 mL DMEM (supplemented with 10% FBS and 1% Penicillin-Streptomycin, 2mM glutamine) were plated in 6-well plate. The following day, 1.5 μg of SrtA plasmid^26^ (pMP71-eGFP-P2A-SrtA-PDGF; provided by Gabriel Victora) and 1.5 μg pECO envelope plasmid were mixed in 250 μl of OptiMEM medium, supplemented with 6 μl FuGENE, and incubated for 20 min. The mix was added dropwise to the BOSC23 cells and incubated at 37°C. 72 hr later, virus supernatant was collected and stored at –80°C for later use.

To generate SrtA-expressing AKP and pro-Met organoids, organoids were dissociated into single cells as described earlier. 2×10^5^ organoid cells resuspended in less than 50 μl ADF^+^ media were mixed with 500 μl of SrtA-virus supernatant described above, 5 μg polybrene, 10 μM Y-27632, and plated in 12-well plate. Cells were spun for 1.5 hr at 2230 rpm, 25°C, then incubated at 37°C incubator overnight. The following day, cells were washed and cultured in matrigel as described above for organoid culture. GFP^+^ SrtA^+^ FLAG^+^ AKP and pro-Met cells were sorted and expanded *in vitro* for future use.

### Generation of AKP.*Ciita*^KO^-organoids

AKP organoids were dissociated to single cells as described above and resuspended in 100 μl OPTI-MEM. Ribonucleoprotein (RNP) complexes were formed by mixing 1.64 μl (0.1 nmol) Alt-R Cas9 (IDT) with 3 μl (0.3 nmol) synthetic sgRNA (IDT) and incubating for 10–20 min at room temperature. Cells were then added to the RNP mix, 100 μl transferred to a 2-mm gap cuvette and electroporated using a NEPA21 electroporator with the following poring pulse parameters: 175 V, 5-ms length, 50-ms interval, two pulses. Electroporated organoids were resuspended gently in pre-warmed minimal medium and incubated at 37 °C for 15 min before plating in Matrigel.

The sgRNAs used for electroporation was:

CIITA sgRNA: AGGUCCUUGAUUAUAUCGUG

### Generation of pro-Met.*Ciita*^OE^-organoids

Pro-Met organoids were dissociated to single cells as described above, and 1×10^5^ organoid cells resuspended in less than 500 μl ADF^+^ media were mixed with 10 uL CIITA virus vector (pLV[Exp]-Puro-EF1A>mCiita [NM_001302618.1], Vector Builder, VB900125-2432nxx), 5 μg polybrene, 10 μM Y-27632, and plated in 12-well plate. Cells were spun for 1.5 hr at 2230 rpm, 25°C, then incubated at 37°C incubator overnight. The following day, cells were washed and cultured in matrigel as described above for organoid culture. MHCII^+^ pro-Met cells were sorted and expanded *in vitro* for future use.

### Colonoscopy-guided tumor injections

Colonoscopy was performed using the Karl Storz colonoscope system composed of Image 1 Spies H3-Z HD Camera (part TH100), Image 1 Connect and H3-Link, CCU modular (parts TC200, TC300), Power LED 300 and light cable (parts TL300 and 495ND), Hopkins Telescope (part 64301AA) ^22,24^. For colonoscopy-guided injection, a custom injection needle (Hamilton, 7803-05), a syringe (Hamilton, 7656-01), a transfer needle (Hamilton, 7770-02), and a colonoscope are assembled in examination sheath (part 61029DK). Mice were anesthetized with 2-5% isoflurane before and during the procedure. To prepare cells for orthotopic organoid transplantations, organoids were first dissociated with cold PBS, washed, then dissociated with TrypLE Express for 30 min at 37 °C, passed through a 70-μm filter, washed, and resuspended in ADF^+^ medium (described under Tumor organoids and cell lines),10% matrigel, 1% filter-sterilized Methylene Blue, and 10 μM Y-27632 supplementation. Cells were administered over 3 injections (7×10^4^ cells in 50 μL per injection), spaced ∼ 0.5 cm apart in the distal colon and rectum. To prepare MC38 and MC38 LVM2 cells for orthotopic injections, cells were dissociated with Trypsin-EDTA (0.25%) for 3 min, washed, and resuspended in in DMEM^+^ medium (described under Tumor organoids and cell lines),10% matrigel, 1% filter-sterilized Methylene Blue, and 10 μM Y-27632 supplementation. Three injections of 2×10^4^ cells per 50 μL were performed per mouse.

### Intrasplenic tumor injections

Mice were anesthetized with 2-5% isoflurane and injected with ketamine for perioperative analgesia at 0.1mg/gram body weight. The abdominal flank fur was shaved, and the skin was disinfected with betadine and 70% ethanol. A small left subcostal incision was made to access the spleen. The spleen was gently exteriorized, and 5×10^5^ AKP or pro-Met organoid cells in PBS (dissociated as described earlier) were injected in the spleen using an insulin syringe. The splenic vasculature and its connection to the pancreas were carefully cauterized using a heat pen, and the spleen was removed from the animal. The pancreas was repositioned within the abdominal cavity. The peritoneum was closed with Polyamide 6 sutures, and the skin incision was closed using clip sutures.

### Histopathology

Representative colon tumors (or entire colons with tumors processed into swiss rolls) of mice orthotopically injected with AKP or pro-Met organoids were harvested 2 and 6-7 weeks post injection, fixed in 4% paraformaldehyde overnight, dehydrated with 50% ethanol followed by 70% ethanol, then cleared in xylene, and paraffin-embedded using a tissue processor (Leica ASP6025, Leica Biosystems, Deer Park, IL). Paraffin blocks were sectioned at 5 μm, stained with hematoxylin and eosin (H&E), and examined by a board-certified veterinary pathologist (S.E.C.). Microscopic changes within the tumors and adjacent intestinal tissue were graded for necrosis, and fibrosis (desmoplasia) on a numerical scale of 0 - 4, where 0 = normal, 1 = minimal (< 9%), 2 = mild (10-24%), 3 = moderate (25–69%), and 4 = severe (> 70%). Epithelial changes at organoid cell implantation site were graded on a scale of 0 - 4, where 0 = normal, 1 = hyperplasia, 2 = dysplasia, 3 = adenoma, and 4 = adenocarcinoma, according to histopathological criteria for colonic tumors in rodents^67,68^. A composite pathology score was generated for each mouse by summing all subcategory scores. H&E and IHC images were acquired using an Olympus VS200 slides scanner (Evident Scientific, Hamburg, Germany) and analyzed with OlyVIA software version 3.4.1 (Olympus, Germany).

### Immunohistochemistry

All colons were evaluated for T cell subsets (CD4 and CD8) and tdTomato using an IHC method previously validated by the MSK’s Laboratory of Comparative Pathology. Briefly, formalin-fixed paraffin-embedded sections were stained using an automated staining platform (Leica Bond RX, Leica Biosystems). After deparaffinization and heat-induced epitope retrieval, large intestinal sections were incubated with either anti-CD4 (dilution 1:250; 14-9766-82, clone 4SM95), anti-CD8a (dilution 1:1000; 14-0808-82, clone 4SM15), or tdTomato (dilution 1:2000; LS-C340696) antibodies. A rabbit anti-rat secondary antibody (for CD4 and CD8 IHC) or a rabbit anti-goat antibody (for tdTomato IHC; BA-4001, BA5000, Vector Laboratories) and a polymer detection system (PK6100, Vector Laboratories) were then applied to the tissues. The 3,3’-diaminobenzidine tetrachloride (DAB) was used as the chromogen, and the sections were counterstained with hematoxylin and examined by light microscopy. Lymphoid tissues (thymus, spleen, and lymph nodes) from naïve mice or skin-expressing tdTomato from genetically engineered mice were used as positive controls.

### Immunofluorescence (IF) staining

Mouse colon was dissected on ice, opened longitudinally, and washed with ice-cold PBS. Swiss rolls of colon tissue were fixed in 4% PFA at 4°C overnight, washed three times with PBS, and transferred to 30% sucrose for at least 16 hours. Tissue was frozen in OCT, cut into 20μm sections on a cryostat, and kept at −20°C until staining. Slide-mounted frozen sections were thawed at room temperature for 20 min, washed twice with PBS for 5 min each, and once for 10 min in 0.1% Triton X-100 in PBS. Sections were blocked for 30 min in blocking buffer (1x BD Perm/Wash Buffer, 1:200 Fc block). Sections were then incubated with conjugated antibodies (1:100 AF488 anti-mouse CD8a, 1:100 AF647 anti-mouse CD3) in blocking buffer for 1 h at 37°C. After three washes in PBS for 5 min each, stained sections were mounted with Fluoromount-G with DAPI and kept at 4°C until imaging.

### Microscopy imaging for IF

IF images (sham mice) were acquired using a Zeiss LSM 980 confocal microscope. Tiled whole-section views were obtained using a 5x/0.16 NA objective at 1024 × 1024 pixel resolution. Regions containing tdTomato aggregates were identified and imaged at higher magnification using a 20x/0.8 NA objective, with tiled Z-stacks collected at 2048 x 2048 pixel resolution. All images were acquired with identical laser power, detector gain, and offset settings across samples. Tile scans were automatically stitched using the Zeiss ZEN software, and Z-stack image series were imported into Fiji (ImageJ) for maximum-intensity projection. Additional Z-stack images (AKP and pro-Met mice) were acquired using an Olympus FV4000 confocal laser scanning microscope with a 20x/0.80 NA objective lens at 1024 × 1024 pixel resolution. Image acquisition and processing were performed using Olympus cellSens/FV4000 software, and Z-stack image series were imported into Fiji (ImageJ) for maximum-intensity projection.

### Tamoxifen administration

Tamoxifen (50 mg/mL) was dissolved in corn oil and 10 % ethanol, shaken at 45°C for 3-4 h, and stored at – 20°C until use. i*Sell*^Tomato^ mice were treated with two doses of Tamoxifen (5 mg/dose) via oral gavage on days –3 and –1 relative to the tumor or sham injection (for early time point analysis), or around days +29, +31 (4 weeks) after tumor injection. *Pou2f3*^CreERT2^ uLIPSTIC mice used for orthotopic tumor transplantation were treated with three doses of Tamoxifen (5 mg/dose, 1 day apart) at least one week before tumor injections.

### Antibody depletion

For CD4^+^ or CD8^+^ T cell depletion, C57BL/6 mice were injected intraperitoneally with 200 µg/mouse anti-CD4 (clone GK1.5), anti-CD8 (clone 2.43), or isotype control (clone LTF-2) depleting antibodies two days before orthotopic AKP tumor injection. Antibody administration was repeated on the day of tumor injection and every 5-7 days until analysis.

### Substrate administration for uLIPSTIC labeling

Biotin–aminohexanoic acid–LPETGS, carboxy-terminal amide, at 95% purity (biotin–LPTEG), was purchased from LifeTein (custom synthesis) and 20 mM stock solutions were prepared in PBS. For uLIPSTIC labelling experiments, six doses of biotin-LPETG substrate were injected intraperitoneally, 20 min a part, over a course of 2 h (8 µmol/1^st^ dose, followed by 2 µmol for each subsequent dose). Mice were sacrificed for flow cytometry analysis 1 h after the last substrate injection.

### Isolation of T cells from colon epithelium and tumors

Colon intraepithelial lymphocytes (IEL) were isolated as previously described^69^. Briefly, colons were harvested and washed in PBS and 1 mM dithiothreitol (DTT) followed by 30 mM EDTA. IEL were recovered from the supernatant of DTT and EDTA washes. Lymphocytes from colon tumors were obtained by digesting the tissue in 6 mL of RPMI supplemented with 2% FBS, 100 μg/ml DNaseI, and 2 mg/ml Collagenase 8 for 45 min at 37°C. Mononuclear cells were isolated by gradient centrifugation using Percoll. Single-cell suspensions were then stained with fluorescently labeled antibodies as described in flow cytometry section below prior to downstream flow cytometry (analysis or sorting).

### Flow cytometry, staining, and gating strategy

Single-cell suspensions were resuspended in PBS + 0.5% BSA + 1mM EDTA (PBE) and incubated with fluorescently labeled antibodies for surface staining for 30min at 4°C. Cells were washed and resuspended in PBE prior to flow cytometry. The following gating strategy was used to examine CD4^+^ or CD8^+^ T cells: single live lymphocytes (based on size and live/dead fixable dye Aqua or NIR stain), CD45^+^, TCRβ^+^, CD8β^−/+^, CD4^+/-^. For analysis of i*Sell*^Tomato^ mice, the following gating strategy was added: CD62L^−^and Tomato^+^ or Tomato^−^. The same gating strategy described above for i*Sell*^Tomato^ mice was used (CD62L^−^and Tomato^+^) for sorting of cells subjected to scRNA-seq. T cells were sorted on a FACSymphony instrument. For uLIPSTIC experiments, the following gating strategy was used to examine tumor cells: single live cells (based on size and live/dead fixable dye NIR stain), EpCAM^+^, CD45^−^, Tomato^+^, FLAG^+^, Biotin^+^, SrtA-GFP^+^.

Intranuclear staining of Foxp3 was conducted using Foxp3 Mouse Regulatory T Cell Staining Kit according to kit instructions. For analysis of cytokine secretion, total mononuclear cells isolated from the colon epithelium or tumors were plated in 96-well plates and incubated at 37°C with 100 ng/mL phorbol 12-myristate 13-acetate (PMA), 200 ng/mL ionomycin, and 2 mM monensin for 4 hr. Intracellular staining for IFN-γ was conducted in Perm/Wash buffer after fixation and permeabilization in Fix/Perm buffer according to kit instructions. Flow cytometry data were acquired on an FACSymphony A5 cytometer and analyzed using FlowJo 10 software package.

### Organoid-T cell co-cultures

T cells isolated from AKP tumors in *Pou2f3*^CreERT2^ uLIPSTIC mice as described above (Isolation of T cells from colon epithelium and tumors) were stained with flow cytometry antibodies as described. T cells isolated from i*Sell*^Tomato^ mice as described above (Isolation of T cells from colon epithelium and tumors) were stained with flow cytometry antibodies as described above. CD62L^+^, CD62L^−^ Tomato^−^, and CD62L^−^Tomato^+^ CD4^+^ and CD8β^+^ T cells were sorted on a BD FACS Aria III Cell Sorter using a 70 µm nozzle at 45 psi. For combined CD4^+^ and CD8β^+^ co-culture, cells were pooled from colon IEL and tumor (from either AKP or pro-Met mice), and cells were pooled from n=4–5 animals per condition per experiment. For T cells isolated from the i*Sell*^Tomato^ and *Pou2f3*^CreERT2^ uLIPSTIC models, 4500 T cells and 4000 T cell, respectively, were co-cultured with 1000 pro-Met organoid cells that were dissociated prior to the co-culture as described above. Cells were co-cultured in final volume of 10 uL in 50% matrigel. Warm TCM supplemented with 10 ng/mL recombinant murine IL-2 (R&D), 20 ng/mL recombinant murine IL-7 (R&D), and 10 ng/mL recombinant murine IL-15/IL-15Rα complex (ThermoFisher) was added to the solidified co-culture drops. On day 5, co-cultures were imaged using a Keyence BZ-X810 microscope in brightfield mode at 2X objective magnification with 1.5X digital zoom. Supernatants were then collected and stored at −80°C for subsequent cytokine analysis by cytometric bead array, performed according to the manufacturer’s instructions.

Following imaging and supernatant collection, co-cultures were dissociated using TrypLE for 30 minutes to facilitate Matrigel and organoid digestion. Samples were further mechanically dissociated by pipetting, washed with PBS, and processed for flow cytometry staining and analysis.

### scRNA-seq library preparation

T cells isolated from i*Sell*^Tomato^ mice as described above (Isolation of T cells from colon epithelium and tumors) were stained with individual Biolegend TotalSeq C antibodies (1:100) to allow for sample-level barcoding. CD62L^−^ Tomato^+^ CD4^+^ or CD8β^+^ were sorted on a BD FACS Aria III Cell Sorter using a 70 µm nozzle at 45 psi. Samples were pooled, assessed for viability, and quickly loaded into a single well of the 10X Genomics Chromium microfluidic device. A single-cell gene expression library was generated following the manufacturer’s protocol for a Chromium Next GEM Single Cell 5’ Reagent Kit v2 (Dual Index), VDJ, with Feature Barcode technology for Cell Surface Protein (to capture the TotalSeq C sample barcodes). Libraries were sequenced on a NovaSeq S1 flow cell.

### scTCR-seq library preparation

Single cells were index-sorted using a FACS Symphony sorter into 96-well plates containing 5μL of lysis buffer (TCL buffer, Qiagen 1031576) supplemented with 1% β-mercaptoethanol and frozen at −80°C prior to RT-PCR. RNA extraction and RT-PCR for TCRβ were performed as previously described^70^. PCR products were barcoded and submitted for MiSeq sequencing^71^ using the TrueSeq Nano kit (Illumina). MiSeq fastq files were demultiplexed, paired-end reads assembled using PANDASEQ^72^, and collapsed using the FASTX toolkit. Demultiplexed reads were assigned to wells by barcode, and resulting FASTA files were aligned and analyzed using IMGT/HighV-QUEST (imgt.org/HighV-QUEST)^73^. Cells sharing identical TCR CDR3 nucleotide sequences were defined as clones. Only in frame junction sequences of TCRβ were included in the analysis.

### scRNA-seq and scTCR-seq data analyses

Raw FASTQ sequencing files were filtered, aligned and the generation of feature barcodes was done using Cell Ranger v7.0.1. Samples were run on the same flowcell using 15 individual feature barcodes to separate TILs and IEL cells from Sham, AKP, and pro-Met conditions. VDJ analysis was run using cellranger VDJ mode with default settings. In total, cellranger recovered 8903 cells with a 7357 median read per cell, with 84% of them presenting VDJ reads. The resulting matrices were loaded into R and analyzed using tidyomics (v1.0) and Seurat R packages (v5.3). Cells exhibiting no or more than one ADT read were excluded from downstream analyses. Next, we removed cells showing more than 10% of total mitochondrial gene expression from the analysis. The expression data was normalized using the SCTransform. Dimensionality reduction was performed via principal component analysis (PCA), and UMAP was applied to visualize the clusters. All clustering possibilities were calculated using FindClusters with Louvain algorithm and were then re-checked by hierarchical relationships and biological significance. Final clustering resolution was decided according to biological difference between T-cell types and cell type annotation was performed using enrichment packages ClusterProfiler (v4.6.2) and fgsea (v1.24.0) and manual curation. Differentially expressed genes were determined using the FindMarkers function of markers of each cluster and between the groups. CD4^+^ and CD8^+^ cells were separated using Cd4 (>0.5 expression) and Cd8b1 (> 1 expression). After the separation, CD4^+^ and CD8^+^ T cells were reprocessed using the aforementioned methods and a total of 1680 CD4^+^ T cells and 1592 CD8^+^ T cells were included in the analysis. Functional enrichment for figures was done using fgsea (v1.24.0) against GO BP (c5.go.bp.v7.5.1.symbols). R-generated plots were produced using ggplot2 (v3.5.2), iSEE visualization package (v2.10.0). The full and the separate rds and the processed data are available on GEO: GSE311111: https://www.ncbi.nlm.nih.gov/geo/query/acc.cgi?acc=GSE311111 (token gxinqgkszzabpiz).

Clonal TCR analysis was performed by processing the filtered_contig_annotations.csv output file from cellranger. Briefly, only cells containing TRA, TRB, TRA_TRB, dual TRA and/or dual TRB contigs were used for statistical analysis. Cells harboring the same V and J genes and CDR3 length, as well as TRA and TRB pairing, were annotated as clonally related. Cells containing only TRA or TRB were also assigned as being part of the same TRA_TRB core if they had an exact CDR3 sequence. Morisita horn overlap index was calculated with mh(data, CI = 0.95, resample = 100) function in the divo v1.0.2 R package. TCR repertoire diversity was analyzed by the Diversity 50 index (D50), which was calculated by ranking the clonotypes by frequency and defined D50 as the proportion of clonotypes needed to reach 50% of cells within each group.

### Expression of TCRs into NFAT-GFP hybridomas

We used the retroviral vector pMSCV-mCD4-PIG TCR-OTII as previously described to transduce the NFAT-GFP 58α−β− hybridoma cells with synthetic TCR constructs^38^. TCR sequences are listed in Supplementary Table 1. pMSCV-mCD4-PIG TCR-OTII, our backbone vector for TCR expression, was provided to Twist Bioscience. Sequences of the TCR α and β genes selected (21 TCRs in total) were obtained from the filtered_contig_annotations.csv output file from scRNA-seq. A target sequence consisting of the TCR α and β genes separated by a self-cleaving P2A peptide was synthesized (Twist Bioscience) and used to replace the OTII TCR gene in the pMSCV-mCD4-PIG plasmid. The Platinum-E retroviral packaging cell line (RV-101) was used for the viral packaging of pMSCV-mCD4-PIG TCR vectors generated by Twist. Platinum-E cells were cultured in a 96-well plate to 60–80% confluency and transfected with pMSCV-mCD4-PIG TCR vectors as a DNA–lipid complex using Lipofectamine 3000 (L3000001) following the manufacturer’s instructions. The culture supernatant containing viral particles was harvested 24 h after transfection. Next, 2.5 × 10^4^ NFAT-GFP cells were centrifuged, resuspended in 200 μl of the virus-containing supernatant supplemented with 10 μg/ml protamine sulfate, and plated in U-bottom 96-well plates. NFAT-GFP cells were spin-transduced by centrifugation at 1,000xg at 32°C for 120 min, pipetted up and down, and cultured overnight at 37 °C in a 5% CO_2_ incubator. Cells were then transferred into a flask and incubated for 1–2 weeks with 10 ml of culture medium (DMEM with 10% FCS, penicillin-streptomycin, and 2 mM L-glutamine) under puromycin selection (1 μg/mL). After selection and propagation, TCR expression in NFAT-GFP cells was confirmed using flow cytometry.

### Preparation of antigen extracts

#### Tumor antigen extracts

AKP and pro-Met *in vitro* organoid lysates were prepared by dissociating organoids into single cells as described above. Cells were resuspended in PBS at 1×10^7^ cells/ mL. Cell suspension was subjected to 5 freeze-thaw cycles in liquid nitrogen and 56°C water bath. Cells were spun down at 1700 g for 5 min to remove cell debris. The supernatant was used a source of tumor lysates, and 5 uL was used for the NFAT-GFP co-culture assay. To prepare AKP *ex vivo* tumor lysate, AKP tumors were collected 2 weeks after orthotopic AKP organoid injections in SPF mice. Tumors were chopped into small pieces and resuspended in PBS supplemented with protease inhibitor cocktail (Roche). Tissues were heated for 1 h at 56°C, followed by 4 freeze-thaw cycles. Tubes were spun down to remove remaining tissue pieces, and the supernatant was used for the NFAT-GFP co-culture assay.

#### Microbial antigen extracts

Total fecal extracts were prepared by dissolving a small fecal pellet in 200 μL of PBS and centrifuged briefly in a benchtop centrifuge for 3-5 seconds to remove large fecal particles. Supernatant was transferred to another tube, centrifuged at 10,000 g for 10 min, then the bacterial pellet was resuspended in 200 μL of PBS. Fecal lysate was heat-inactivated at 75°C for 1 h. To prepare bacterial antigen extracts from MM12 species, bacteria recovered from glycerol stocks were grown anaerobically at 37°C for 1-2 days in liquid AAM^+^ media (containing 18 g/L BHI, 15 g/L trypticase soy broth, 5 g/L yeast extract, 2.5 g/L K2HPO4, 0.5 g/L d-glucose, 0.25 g/L cysteine-HCl.H2O, 0.25 g/L Na2S.9H2O, 1 mg/L hemin, 0.5 mg/L Vitamin K3, 3% heat-inactivated FCS). Bacteria were then plated on Columbia blood agar (CBA) plates and (anaerobe systems) grown anaerobically for 48-72 hr. Single colonies were isolated and grown in AAM^+^ liquid media overnight. 2 mL of liquid media was spun down at 10,000 g for 10 min in sterile Eppendorf tubes and resuspended in 1 mL sterile PBS and heat-inactivated at 75°C for 1 h prior to use. *Helicobacter hepatics* was provided by Dan Littman (NYU) and grown as previously described^74^. Optical density (OD) of fecal and bacterial lysates was measured using nanodrop and normalized to OD ≤ 1, and 1 μL of fecal or bacterial lysates was used per well in the *in vitro* coculture assay described below.

#### Dietary antigen extracts

Food and dietary protein-derived antigen mixtures (for casein, soy, and zein) were prepared as described previously^75^. Briefly, solid food pellets were ground using pestle, and 50 mg of powder was resuspended in 1mL of PBS and 50 μg of 50 mg/ mL stock was used per well in the *in vitro* coculture assay.

### *In vitro* screening for TCR specificity

At least seven days prior to the NFAT-GFP reporter assay, CD45.1^+^ C57BL/6 mice were injected subcutaneously with 1×10^6^ B16-Flt3L melanoma cells^38,76^ to enrich for dendritic cells (DCs) in the spleen. Tumor were harvested no more than 4 weeks post-injections. On the day of the experiment, tumor-injected CD45.1 mouse spleens were harvested and digested in 400 U/mL collagenase D and 10 µM EDTA. Red blood cells were lysed using Red Blood Cell Lysis Buffer. CD11c^+^ dendritic cells were enriched using mouse CD11c MicroBeads UltraPure.

For the NFAT-GFP co-culture assay, 5×10^4^ splenic DCs were cultured in 100 µL in a 96-well U-bottom plate and pulsed with antigens of interest for 2-4 hr at 37 °C and 5% CO_2_. Antigens were obtained as described above. Next, 1×10^4^ TCR-transduced NFAT-GFP cells were cocultured in a 96-well plate with the pre-pulsed DCs. After 18–40 hr, cells were stained for Live/Dead Zombie NIR, TCRβ, CD4 and CD45.1, and GFP expression was measured by flow cytometry. As positive control, TCR responsiveness of each NFAT-GFP hybridoma line was determined by anti-CD3 stimulation (0.5 µg/well).

### Whole Exome Sequencing (WES) of tumors

AKP and pro-Met tumor organoids (1×10^6^ each, frozen pellets) were submitted to GENEWIZ (Azenta Life Sciences, South Plainfield, NJ) for whole-exome sequencing. Sequencing libraries were prepared using the Twist Mouse Exome Panel and sequenced with paired-end 150-bp reads on an Illumina platform. Raw sequencing data were processed and analyzed using the GENEWIZ bioinformatics pipeline. Briefly, raw reads were trimmed with Trimmomatic (v0.39), aligned to the Mus musculus GRCm38 reference genome using Sentieon (202308), and processed for duplicate marking. Somatic single-nucleotide variants (SNVs) and small insertions/deletions (indels) were identified using the Sentieon TNseq algorithm. Variants were normalized with bcftools (v1.13) and annotated using Ensembl Variant Effect Predictor (VEP, release 104). An annotated joint somatic mutation annotation format (MAF) file generated by GENEWIZ was imported into R and analyzed using the maftools package. Genes uniquely mutated in the AKP or pro-Met tumors were identified by comparing gene-level mutation profiles between samples, and mutation frequencies were compared using the mafCompare function.

### Statistical analyses

Statistical analyses were performed using GraphPad Prism v.10. Flow cytometry analysis was performed using FlowJo software. Data in graphs show mean and p values <0.05 were considered to be significant. Adobe Illustrator was used to assemble and edit the figures. R Studio Version 2023.06.0+421 and R version 4.3.2 were used to analyze scRNA-seq data.

---

**Figure S1:**
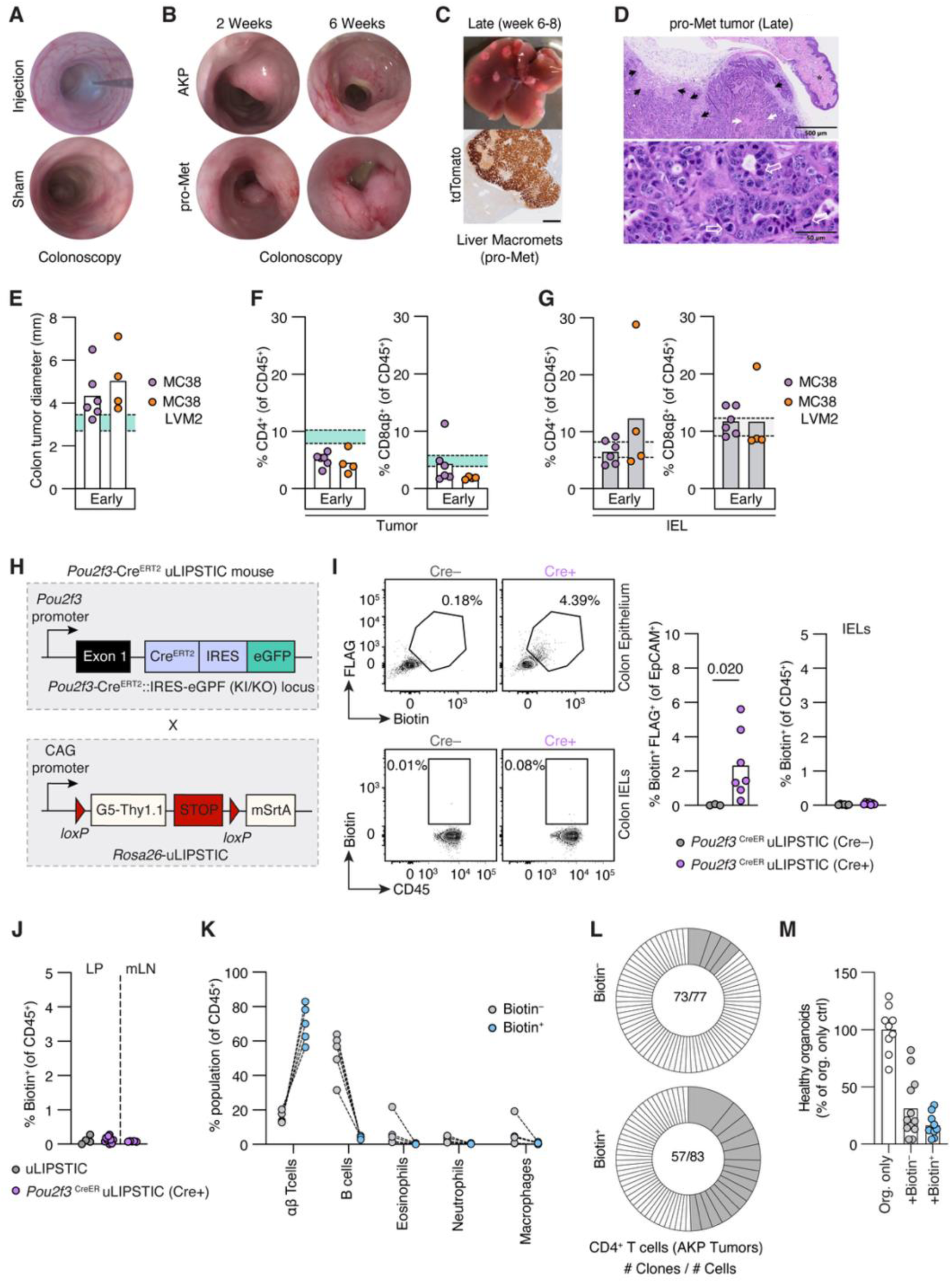
Extended data related to Figures 1 and 2. (A) Colonoscopy images of injection with needle in view (top) and sham control two weeks post-injection (bottom). (B) Colonoscopy images of tumor growth in AKP and pro-Met CRC models at 2 and 6 weeks post-tumor injection. (C) Liver macrometastases (top) and tdTomato immunohistochemistry (bottom) from liver sections in the pro-Met CRC model at 6-8 weeks post tumor injection in the colon. Scale bar = 500 μm. (D) Representative H&E-stained section of a late-stage pro-Met adenocarcinoma expanding into the underlying connective tissue of the rectum (asterisk) and distal colon. The adenocarcinoma (top image) exhibited multifocal areas of necrosis (white arrows) and desmoplasia (reactive fibrous connective; black arrows). High magnification (bottom image) showed the adenocarcinoma is composed by neoplastic columnar to polygonal epithelial cells with small to medium eosinophilic cytoplasm and round to oval nuclei containing one to two prominent nucleoli. Mitotic figures are present (open white arrows). Scale bar = 500 μm for top image, and = 50 μm for bottom image. (E) Colon tumor diameter 2 weeks after the MC38 and MC38 LVM2 tumor injections. Each symbol represents a mouse. Shaded area represents the 95% confidence interval of colon tumor diameter of AKP tumors at the early (2-week) stage. (F) Frequencies of CD4^+^ and CD8αβ^+^ T cells among CD45^+^ T cells in colon tumors of respective tumor lines at the early CRC stage. Shaded area represents the 95% confidence interval of T cell populations in AKP tumors at the early stage. (G) Frequencies of CD4^+^ and CD8αβ^+^ T cells among CD45^+^ IEL isolated from colon epithelium of tumor-injected mice at the early CRC stage. Shaded area represents the 95% confidence interval of T cell populations in Sham IEL. (H) Schematic diagram showing the genetics of the *Pou2f3*^CreERT2^ uLIPSTIC model. (I) *Pou2f3*^CreER^ uLIPSTIC mice (either Cre– or Cre+) were treated by oral gavage with 200 µL of 50 mg/mL tamoxifen on days −3 and −1. On day 0, six doses of biotin-LPETG substrate were injected intraperitoneally over a course of 2 h (8 µmol/1^st^ dose, followed by 2 µmol for each subsequent dose). Mice were sacrificed for flow cytometry analysis 1 h after the last substrate injection. Dot plots and frequency of FLAG and biotin expression in colonic epithelial cells (top). Dot plots and frequency of biotin expression in CD45^+^ cells harvested from colon epithelium (bottom). (J) Frequency of biotin^+^ among CD45^+^ cells isolated from colon lamina propria (LP) and mesenteric lymph nodes (mLN) of AKP^SrtA^-injected mice following biotin-LPETG substrate administration. (K) Frequencies of indicated immune cell populations among biotin^−^ and biotin^+^ CD45^+^ cells. (L) Pie charts showing biotin^−^ (top) and biotin^+^ (bottom) CD4^+^ T cell clones isolated from AKP^SrtA^ tumors two weeks post transplantation in *Pou2f3*^CreER^ uLIPSTIC mouse. Grey sections represent expanded clones (detected more than twice), and white sections represent singlet clones. Number inside chart represents the number of CD4^+^ T cell clones / number of recovered CD4^+^ T cells. Each pie chart represents one mouse. (M) Healthy organoid frequency calculated by normalizing the number of healthy organoids recovered per well to the average number of healthy organoids recovered in control (organoids only).

**Figure S2:**
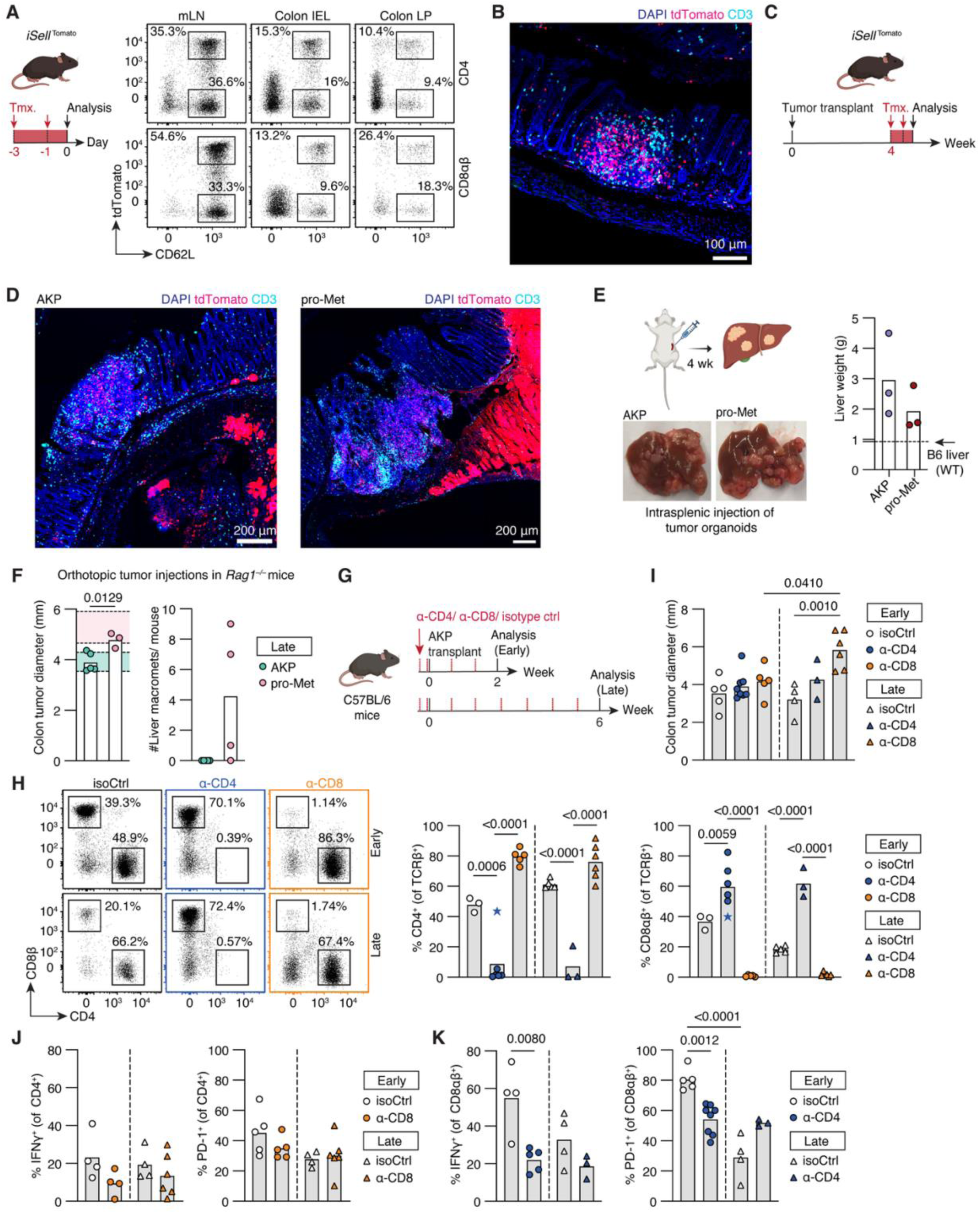
Effects of modulating the adaptive immune system on AKP tumor outcome, and extended controls for Figure 3. (A) Experimental design (left) and dot plot (right) showing CD62L^+^ Tomato^+^ and Tomato^−^ among CD4^+^ (top) and CD8αβ^+^ (bottom) T cells across the mLN, colon IEL and LP 1 day after the second dose of tamoxifen administration in i*Sell*^Tomato^ mice. (B) Immunofluorescence (IF) staining of mouse colon harvested 1 day after the second dose of tamoxifen administration in i*Sell*^Tomato^ mice. Staining shows DAPI, tdTomato, and CD3. Magnification= 20X. Images are representative of n=2 mice. (C) Experimental design. i*Sell*^Tomato^ mice were transplanted with AKP or pro-Met organoids, and treated with tamoxifen 4 weeks later. Colons were isolated for IF staining one day after the second tamoxifen dose. (D) IF staining of colons isolated from (C). tdTomato represents both tumor cells (bright on the right of the images), and also tdTomato^+^ immune cells. Images are representative of n=2 mice/ condition. (E) Scheme of the intrasplenic injection model and display of AKP and pro-Met liver tumor growth. Quantification of tumor load by liver weight. The dashed line indicates average healthy adult B6 liver weight^77^. (F) *Rag1^−/–^* mice were orthotopically injected with AKP or pro-Met tumor organoids using a colonoscope. Average colon tumor diameter per mouse (left) and number of liver macrometastases (right) were assessed 6-7 weeks after tumor injection. Shaded areas represent the 95% confidence interval of colon tumor size in AKP (green) and pro-Met (pink) tumor organoids injected into immunocompetent SPF mice. (G) Experimental design for panels H–K. C57BL/6 mice were injected with 200 µg/mouse anti-CD4 (α-CD4), anti-CD8 (α-CD8), or isotype control (isoCtrl) two days before orthotopic AKP tumor organoid injection. Antibody injections were repeated on the day of tumor injection and every 5-7 days. The dashed red line indicates antibody injections. Mice were sacrificed for analysis 2 or 6 weeks after tumor injection. (H) Dot plots and frequencies of CD4^+^ (middle) and CD8β^+^ (right) among TCRβ^+^ T cells at the indicated conditions and time points. The blue star symbol refers to a mouse with unsuccessful CD4 T cell depletion and hence was excluded from further analysis. (I) Average colon tumor diameter per mouse under the indicated treatment conditions and analysis time points. (J) Frequency of IFN-γ^+^ (left) and PD-1^+^ (right) among CD4^+^ T cells isolated from colon tumors following *ex vivo* PMA/ionomycin stimulation. (K) Frequency of IFN-γ^+^ (left) and PD-1^+^ (right) among CD8αβ^+^ T cells isolated from colon tumors following *ex vivo* PMA/ionomycin stimulation. Unpaired Student’s t-tests were performed for (E) and (F). Ordinary one-way ANOVA with Tukey’s multiple comparison tests were performed in (I–K).

**Figure S3:**
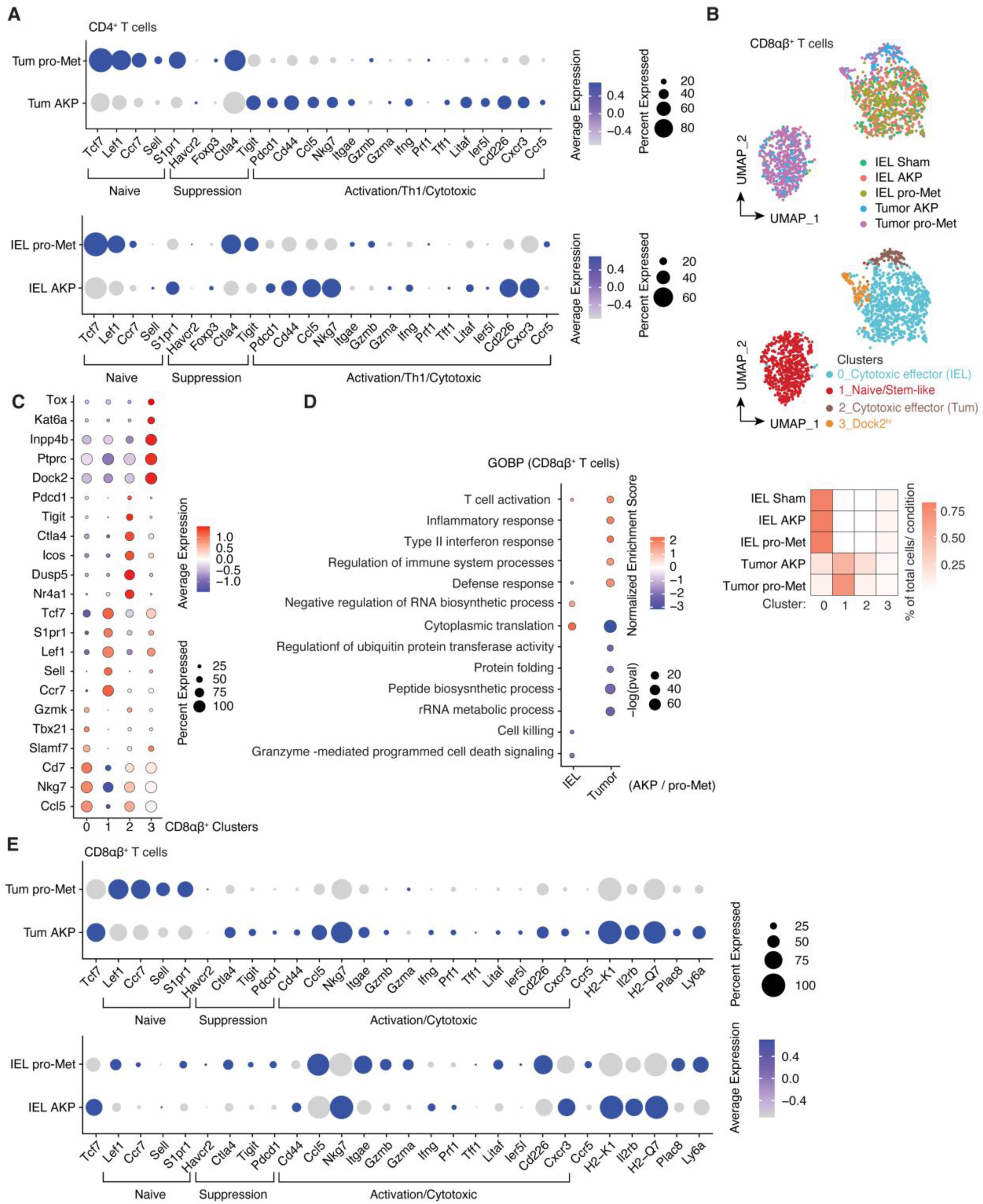
Transcriptional profiles of recently recruited CD4^+^ and CD8αβ^+^ T cells during the early CRC stages. Extended data for Figure 4. i*Sell*^Tomato^ mice orthotopically injected with sham, AKP, or pro-Met tumors using colonoscope 1 day post second tamoxifen dose. CD62L^−^ Tomato+ TCRαβ^+^ T cells from colon IEL and tumor were single-cell RNA sequenced. CD4^+^ and CD8αβ^+^ T cells were separated for further analysis. (A) Dot plot of selected differentially expressed genes between AKP and pro-Met tumor-derived CD4^+^ T cells (top) or AKP IEL and pro-Met IEL-derived CD4^+^ T cells (bottom). (B) UMAP of CD8αβ^+^ T cells visualizing cells isolated per condition (top), main identified clusters using Seurat (SCT normalization, resolution = 0.3) (middle), and a heat map (bottom) showing the frequency of cells per condition across clusters. Each condition (row) represents 100%. (C) Dot plot of CD8αβ^+^ T cell clusters visualized in (B), showing the selected markers expressed per cluster. (D) Selected Gene Ontology Biological Processes (GOBP) differentially expressed between AKP, pro-Met IEL and tumor-derived CD8αβ^+^ T cells. Colors represent the Normalized Enrichment Score (NES) associated with that pathway in IEL and Tumor and the dot radius represents -log of the adjusted p-value of that enrichment. (E) Dot plot of selected differentially expressed genes between AKP and pro-Met tumor-derived CD8αβ^+^ T cells (top) or AKP IEL and pro-Met IEL-derived CD8αβ^+^ T cells (bottom).

**Figure S4:**
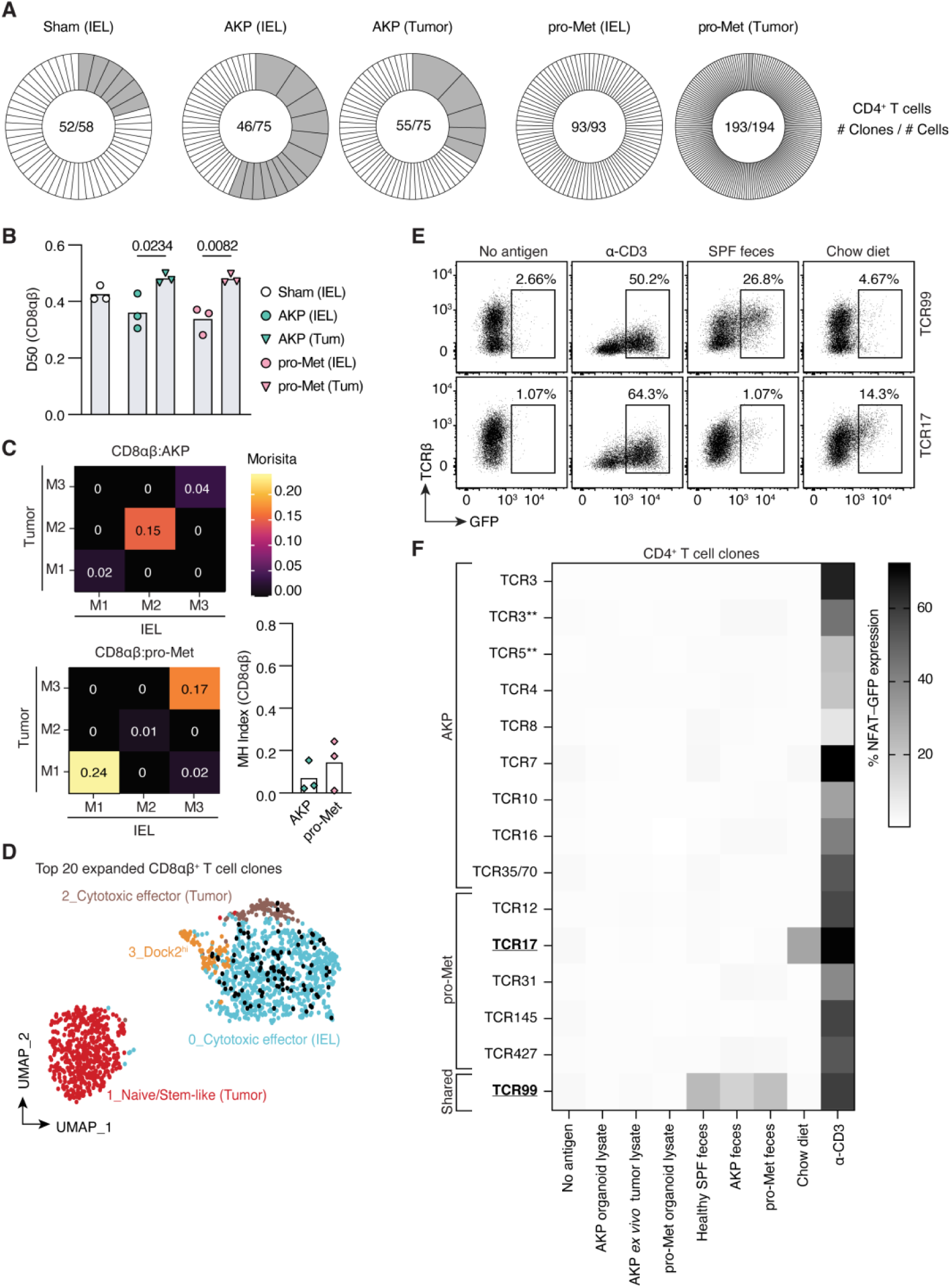
Minimal clonal expansion and clonal overlap of CD8αβ^+^ T cells across conditions, and extended data related to Figure 6. (A) Pie charts showing CD62L^−^ Tomato^+^ CD4^+^ T cell clones isolated from i*Sell*^Tomato^ mice (IEL and Tumor) two weeks post tumor transplantation. Grey sections represent expanded clones (detected more than twice), and white sections represent singlet clones. Number inside chart represents the number of CD4^+^ T cell clones / number of recovered CD4^+^ T cells. Each pie chart represents one mouse. (B) Diversity index 50 (D50) of CD8αβ^+^ T cell clones isolated from the indicated tissues and conditions. (C) CD8αβ^+^ T cell clonal sharing between IEL and tumor in AKP (top) and pro-Met (bottom) injected mice, represented by the MH index. M1,2,3 = Mouse 1,2,3. (D) Top 20 expanded CD8αβ^+^ T cell clones (black) overlayed on CD8αβ UMAP. Seurat-identified clusters (resolution =0.3) were visualized and labelled on the UMAP. (E–F) TCRs from expanded CD4^+^ T cell clones were reconstructed and expressed in TCR-deficient murine NFAT-GFP T cell hybridomas and co-cultured with antigen-pulsed splenic dendritic cells (DCs) for 18–40 h. NFAT-GFP expression indicating TCR reactivity is represented by dot plots in (E) and summarized in the heatmap in (F) for cloned NFAT hybridoma cell lines with high TCRβ expression. Frequencies indicate GFP expression out of CD4^+^ T cells in response to no antigen stimulation, anti-CD3 treatment, or stimulation with tumor, fecal, or dietary proteins. AKP and pro-Met organoid lysates were prepared from *in vitro* cultured organoids. AKP *ex vivo* tumor lysate was prepared from AKP tumors collected 2 weeks after orthotopic tumor injections. AKP and pro-Met mouse fecal lysates were collected from early-stage AKP or pro-Met injected mice. Cloned TCR sequences are listed in Supplementary Table 1. Underlined TCR clones indicate clones with detected reactivities to broad proteins screened in this panel. Ordinary one-way ANOVA with Tukey’s multiple comparison test was performed in (B).

**Figure S5:**
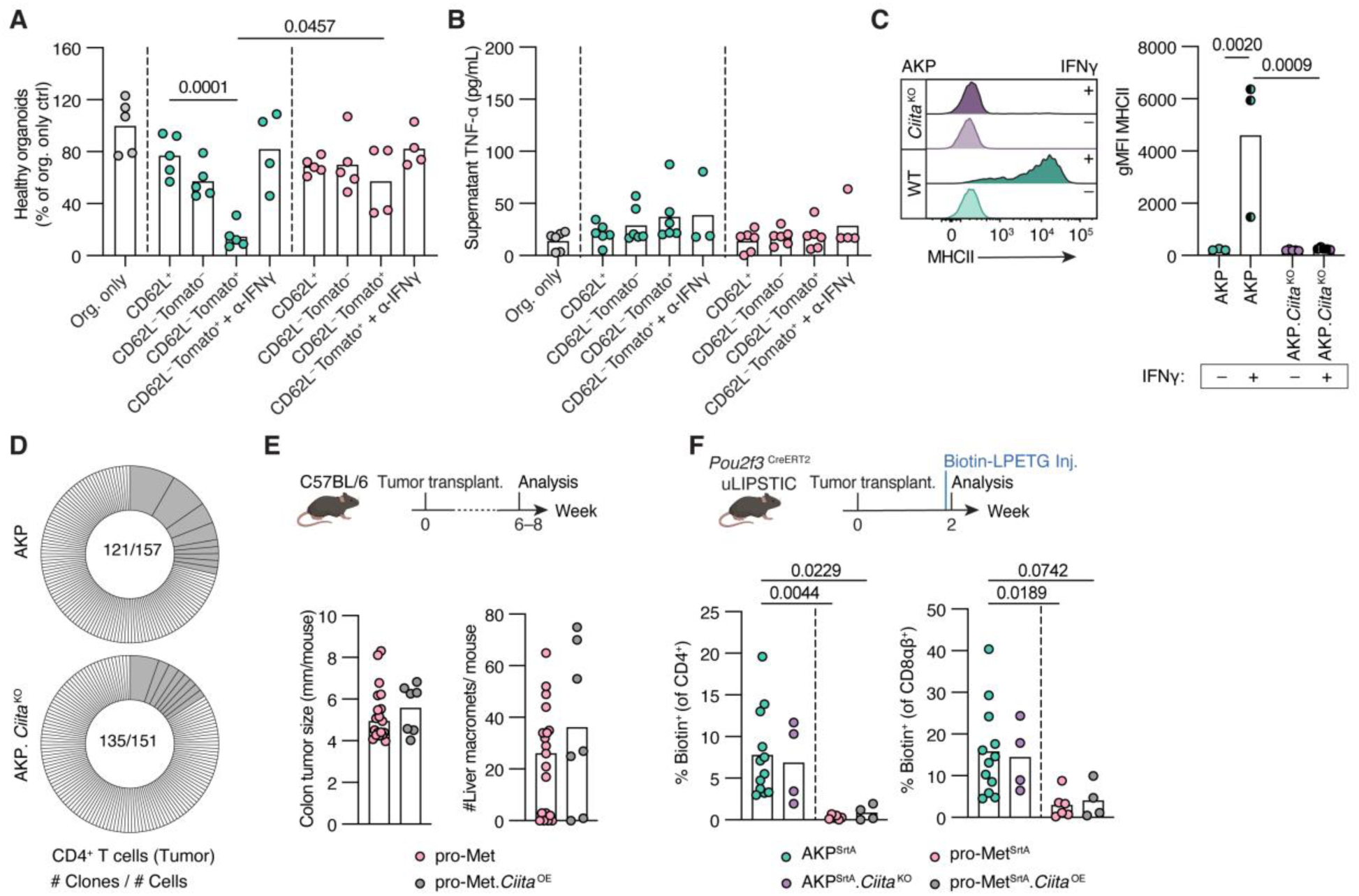
Extended data related to Figures 7 and 8. (A) Healthy organoid frequency calculated by normalizing the number of healthy organoids recovered per well to the average number of healthy organoids recovered in control (organoids only). (B) CBA analysis of co-culture supernatant collected on day 5. Graph shows levels of TNF-α per condition. (C) MHCI gMFI of AKP and AKP.*Ciita* ^KO^ organoids in the presence or absence of IFN-γ stimulation. (D) Pie charts showing CD62L^−^ Tomato^+^ CD4^+^ T cell clones isolated from i*Sell*^Tomato^ mice (IEL and Tumor) two weeks post AKP and AKP.*Ciita* ^KO^ tumor transplantation. Grey sections represent expanded clones (detected more than twice), and white sections represent clones detected once. Number inside chart represents the number of CD4^+^ T cell clones / number of recovered CD4^+^ T cells. Each pie chart represents one mouse. (E) Average colon tumor size per mouse (left), and liver macrometastases (right) at the late tumor stage (6-8 weeks post transplantation). Pro-Met control shown in (E) is from the same dataset in Figures 1B and 1C. Ordinary one-way ANOVA with Tukey’s multiple comparison test was performed for colon tumor size with all groups combined. (F) Frequencies of biotin^+^ among CD4^+^ (left) and CD8αβ^+^ (right) T cells harvested from the indicated tumors. Data from AKP^SrtA^ and pro-Met^SrtA^ is the same as in Figure 2C. Ordinary one-way ANOVA with Tukey’s multiple comparison test was performed with all groups combined.

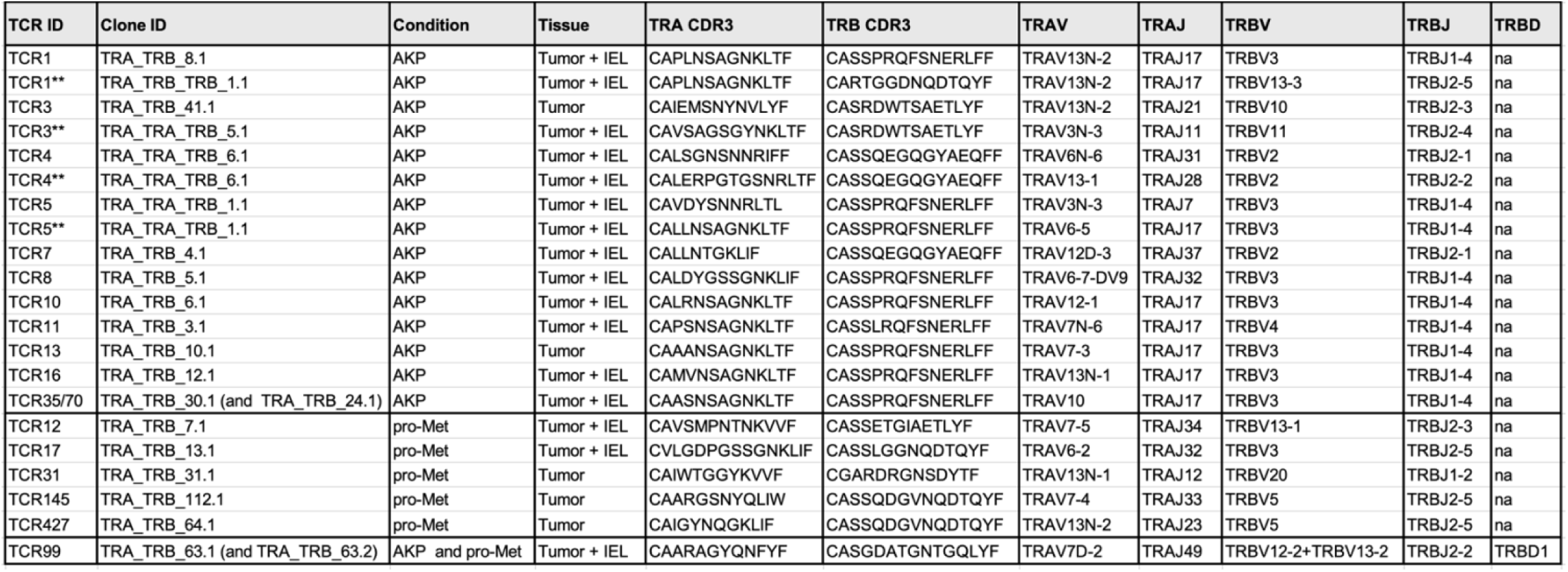

## Declaration of generative AI and AI-assisted technologies in the manuscript preparation process

During the preparation of this work the authors used ChatGPT for grammatical corrections. After using this tool/service, the authors reviewed and edited the content as needed and take full responsibility for the content of the published article.

